# Structural Basis for Iterative Methylation by a Cobalamin-dependent Radical S-Adenosylmethionine Enzyme in Cystobactamids Biosynthesis

**DOI:** 10.1101/2025.11.17.688841

**Authors:** Jiayuan Cui, Bo Wang, Ravi K. Maurya, Squire J. Booker

## Abstract

Cystobactamids are non-ribosomal peptide natural products that function as DNA gyrase inhibitors, exhibiting significant antibacterial activity. They are isolated from Cystobacter sp. Cbv34 and contain various alkoxy groups on para-aminobenzoic acid moieties, which are believed to play a crucial role in antibacterial functions. The alkoxy groups are generated by iterative methylations on a methoxy group by the cobalamin (Cbl)-dependent radical S-adenosylmethionine (SAM) enzyme CysS. CysS catalyzes up to three methylations to give ethoxy, isopropoxy, sec-butoxy, and tert-butoxy groups. For each methylation, CysS uses a ping-pong mechanism in which two molecules of SAM are consumed. One SAM is used to methylate cob(I)alamin, while another generates a 5′-deoxyadenosyl 5′-radical to initiate substrate methylation. However, little is known about how the enzyme promotes both Cbl methylation and iterative substrate methylation, which occur by polar S_N_2 and radical processes, respectively. Here, we report three X-ray crystal structures of a homolog of CysS from *Corallococcus sp. CA054B*. Two were determined in the presence of methoxy- and ethoxy-containing substrates, showing how CysS accommodates substrates and products during iterative methylation. The third structure, determined in the absence of a substrate, exhibits structural changes that reorient the SAM’s conformation to allow for the methylation of cob(I)alamin.

## Introduction

Cystobactamids are a class of nonribosomal peptide natural products isolated from *Cystobacter sp. Cbv34* that inhibit the growth of various pathogens belonging to the “ESKAPE” panel (1, 2). This panel includes Gram-positive and Gram-negative organisms resistant to multiple antibiotics, and they are the leading cause of hospital-associated infections (3). Cystobactamids are DNA gyrase inhibitors that do not possess significant cross-resistance with fluoroquinolones, the most important clinically used gyrase inhibitors (1, 4, 5). In addition, some cystobactamid derivatives are not exported by major efflux pumps, a significant mechanism of antibiotic resistance(4). The chemical structures of cystobactamids include a β-methoxyasparagine hinge moiety and several *para*-aminobenzoic acid (PABA) units that extend from both the carboxyl and amino groups of the hinge moiety *via* amide bonds (**Figure 1A**) (1, 6–8). Intriguingly, cystobactamids exist as a panel of analogs containing methoxy (OMe), ethoxy (OEt), isopropoxy (O*^i^*Pr), and sec-butoxy (O*^s^*Bu) groups on the PABA units (5, 9, 10). Structure-activity relationship studies have established the importance of the alkoxy groups on the PABA moieties for antibacterial activity (5, 9–12).

**Figure 1.**
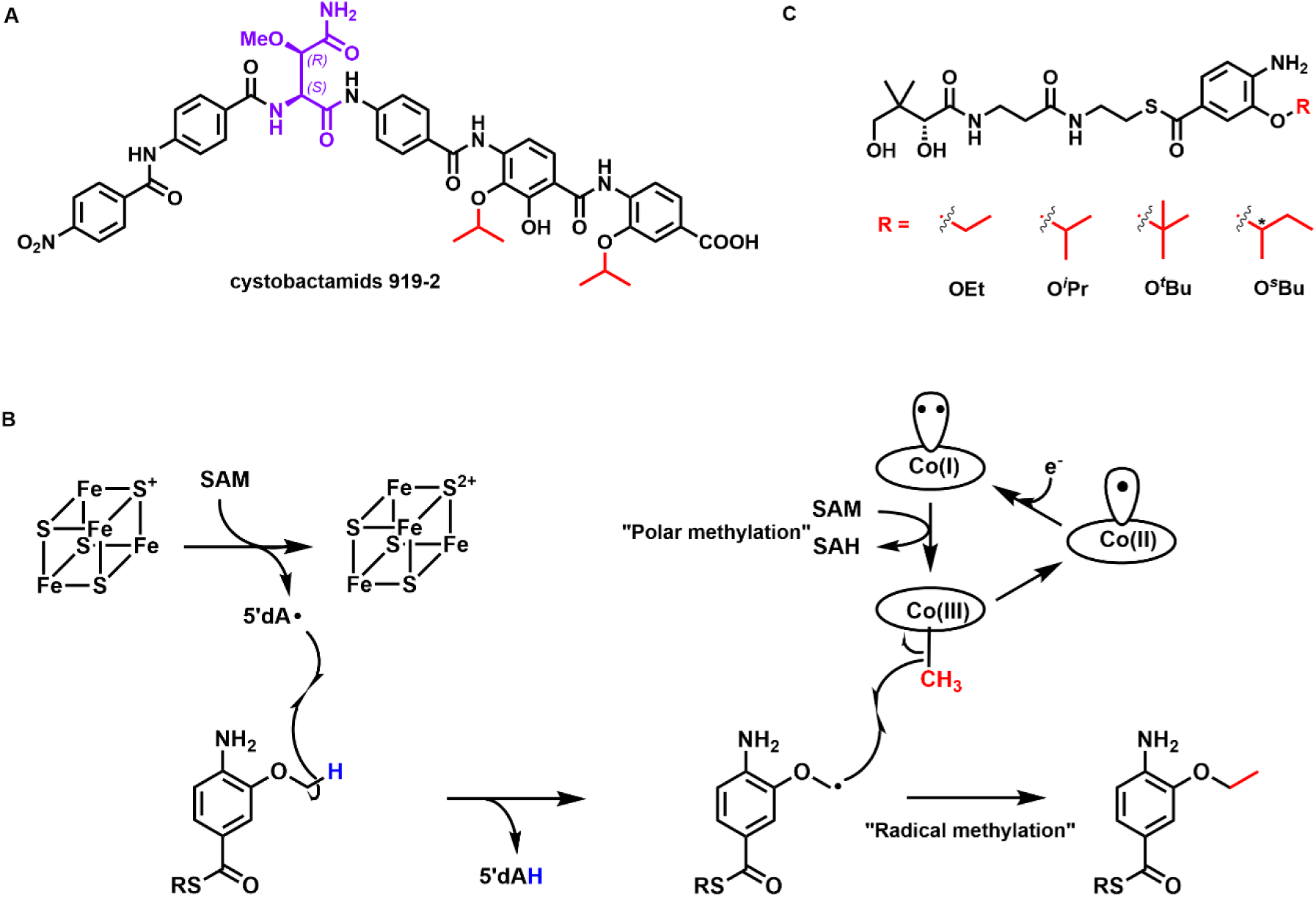
**A.** The chemical structure of cystobactamids 919-2 isolated from *Cystobacter sp. Cbv34*. The stereochemistry of the hinge moiety was initially assigned to be (*2S, 3S*) (1), which was later revised to be (*2S, 3R*) by total synthesis (7–8); **B.** Proposed catalytic mechanism of CysS; **C.** The CysS reaction with a pantetheinylated 3-methoxy-4-amino-benzoic acid substrate gives various methylated products, including OEt-, O*^i^*Pr-, O*^t^*Bu-, and O*^s^*Bu-containing species. The asterisk in O*^s^*Bu indicates that the stereochemistry was not determined.

The biosynthesis of cystobactamids has been reconstituted *in vivo* and *in vitro* (1, 13). The current biosynthetic model requires a six-module assembly line encoded by two non-ribosomal peptide synthetases (NRPS), CysK and CysG. Each module incorporates a β-methoxyasparagine, a PABA, or a modified PABA into the structure of cystobactamids (**Figure S1A**). The tailoring of a PABA by three enzymes, CysC, CysF, and CysS, occurs before it is incorporated into the cystobactamids’ structure (**Figure S1B**). CysC is an oxygenase that adds hydroxyl groups to PABA’s 3-position, or both 2- and 3-positions. CysF is an *S*-adenosylmethionine (SAM)-dependent methyltransferase that methylates the 3-OH group of PABA. The final tailoring step is the iterative methylation on the resulting 3-methoxy group of a PABA by CysS, a cobalamin (Cbl)-dependent radical SAM (RS) enzyme, to yield ethoxy, isopropoxy, tert-butoxy (O*^t^*Bu), or sec-butoxy groups, all of which are observed in isolated cystobactamids. The finding that cystobactamids can be isolated with various alkoxy groups suggests that the entire biosynthetic process tolerates modifications to the alkoxy moiety, providing a strategy to engineer analogs with improved antibacterial properties.

Currently numbering over 800,000 individual sequences, RS enzymes constitute one of the largest enzyme superfamilies (14–19). Each contains a [Fe_4_S_4_] cluster coordinated by three Cys residues, overwhelmingly found in a CX_3_CX_2_C motif. The iron-sulfur (FeS) cluster is required to cleave SAM reductively to methionine and a 5′-deoxyadenosyl 5′-radical (5′dA•) (14, 15). The role of the 5′dA• is to abstract a hydrogen atom (H•) from a bound substrate, a step that initiates more than 100 distinct reaction types (14–19). CysS belongs to a large subfamily of RS enzymes that require a Cbl cofactor in addition to the FeS cluster (20–22). Most of these enzymes are methylases that use both cofactors to methylate sp^2^- and sp^3^-hybridized carbons or phosphinate phosphorus centers, often found in medicinally important natural products and some proteins (19).

Wang *et al*. showed that CysS methylates the methoxy group of a pantetheinylated 3-methoxy-4-amino-benzoic acid group using a ping-pong mechanism (**Figure 1B**) (23–25). The reaction is initiated by a polar transfer of a methyl group from SAM to cob(I)alamin, yielding methylcobalamin (MeCbl) and *S*-adenosylhomocysteine (SAH). After dissociation of SAH, a second SAM molecule binds and undergoes a reductive cleavage to give a 5’dA•, which abstracts a H• from the methoxy group. The resulting substrate radical attacks the methyl group of MeCbl, leading to the homolytic cleavage of the Co-Me bond and producing the ethoxy product and cob(II)alamin. Cob(II)alamin is reduced by an external electron to cob(I)alamin, which is then remethylated to MeCbl. A similar mechanism has been proposed for several other cobalamin-dependent RS methylases (Cbl-RSM), except for TsrM (26–29). However, the details of how the enzymes regulate the two distinct SAM-dependent steps are unclear.

CysS is one of a few known Cbl-RSMs that perform iterative methylations, allowing the enzyme to generate a library of products with various alkoxy functionalities (**Figure 1C**) (23, 24). It should be noted that cystobactamids with an O*^t^*Bu group have not yet been isolated from native organisms; however, *in vitro* studies have demonstrated the possible existence of such analogs (23). Only a handful of Cbl-dependent RS enzymes have been characterized structurally: OxsB, TsrM, TokK, Mmp10, and two unannotated DUF512-containing proteins (27, 30–33). Like CysS, TokK also catalyzes iterative methylations, performing three sequential methyl transfers at C6 of a carbapenam substrate, yielding the isopropyl substituent in the carbapenem antibiotic asparenomycin A (22, 27). However, only two structures of TokK have been reported. One contains a bound unmethylated substrate, while the second contains no substrate.

Here, we report three X-ray structures of CysS: one with the OMe substrate, one with the OEt substrate, and one without a substrate. The two substrate-bound structures reveal that CysS interacts with each substrate in a similar manner. Although crystallization with an O*^i^*Pr substrate was unsuccessful, the other structures identified a residue (Phe485) that may play a role in the third methylation. Substitutions at Phe485 alter the catalytic rate and the ratio of products formed from the O*^i^*Pr substrate. In the structure without a substrate, we observe changes in the positions of several residues—including Trp271 and Phe485, as well as a flexible loop near Phe485—compared to the structures containing substrates. Subsequent biochemical studies suggest that these movements facilitate cob(I)alamin methylation during catalysis, addressing a longstanding conundrum in the field.

## Results

### CysS and its homolog catalyze iterative methylation

Our initial structural studies focused on CysS from *Cystobacter sp. Cbv34* (*cb*CysS, UniProt ID: A0A0H4NV78), previously isolated, characterized, and studied mechanistically (23–25). However, we were unable to obtain suitably diffracting crystals of this protein. Therefore, we focused on a homolog from *Corallococcus sp. CA054B* (*cc*CysS, UniProt ID: A0A3A8HCN5), which shares 89% sequence identity with *cb*CysS. Genes for both enzymes were cloned into a SUMO vector to produce an N-terminal SUMO-tagged protein, where the SUMO tag is removed by treatment with the Ulp1 protease. Both proteins were purified to ≥95% homogeneity (**Figure S2**). Iron analysis shows 3.3 Fe/*cc*CysS and 3.7 Fe/*cb*CysS. Both proteins exhibit UV-vis features consistent with the presence of FeS clusters (broad feature at 410 nm) and Cbl (features at 320 nm) (**Figure S3**). The quantification of Cbl shows 0.94 (*cc*CysS) and 0.81 (*cb*CysS) equiv per protein(34). Genes for *cc*CysS and *cb*CysS were co-expressed with the *btu* operon, which encodes proteins involved in Cbl uptake and trafficking (35). Analysis of the purified proteins by mass spectrometry, according to a previously described procedure, shows they contain a mixture of hydroxocobalamin (OHCbl) (35.7% for *cb*CysS; 37.6% for *cc*CysS), MeCbl (33.2% for *cb*CysS; 41.5% for *cc*CysS), and adenosylcobalamin (AdoCbl) (31.1% for *cb*CysS; 20.9% for *cc*CysS)(34).

To determine whether *cc*CysS catalyzes the same reaction as *cb*CysS, we synthesized all five substrates and products (OMe, OEt, O*^i^*Pr, O*^s^*Bu, and O*^t^*Bu) and used them in reactions for both proteins, as previously described (36). It should be noted that the synthesized 3-alkoxy-4-aminobenzoic acid substrates were linked to a pantothenic acid moiety *via* an ester bond (**Figure 2A**) instead of the natively found thioester bond (**Figure 1C**) to avoid potential thioester hydrolysis or transesterification during crystallization (23, 37). We observe that *cc*CysS catalyzes iterative methylations on the synthetic substrates with a catalytic profile similar to that of *cb*CysS. In reactions using the OMe substrate, only the OEt and O*^i^*Pr products are detected (**Figure 2B & E**). In the reaction using the OEt substrate, only the O*^i^*Pr product is detected (**Figure 2C & F**). The O*^s^*Bu and O*^t^*Bu products are observed only when starting from the O*^i^*Pr substrate (**Figure 2D & G**), suggesting that the third methylation is the slowest. Interestingly, *cb*CysS produces the O*^s^*Bu and O*^t^*Bu products in a 0.8:1 ratio, whereas the *cc*CysS reaction produces them in a 2:1 ratio, suggesting that differences in their structures govern product partitioning.

**Figure 2.**
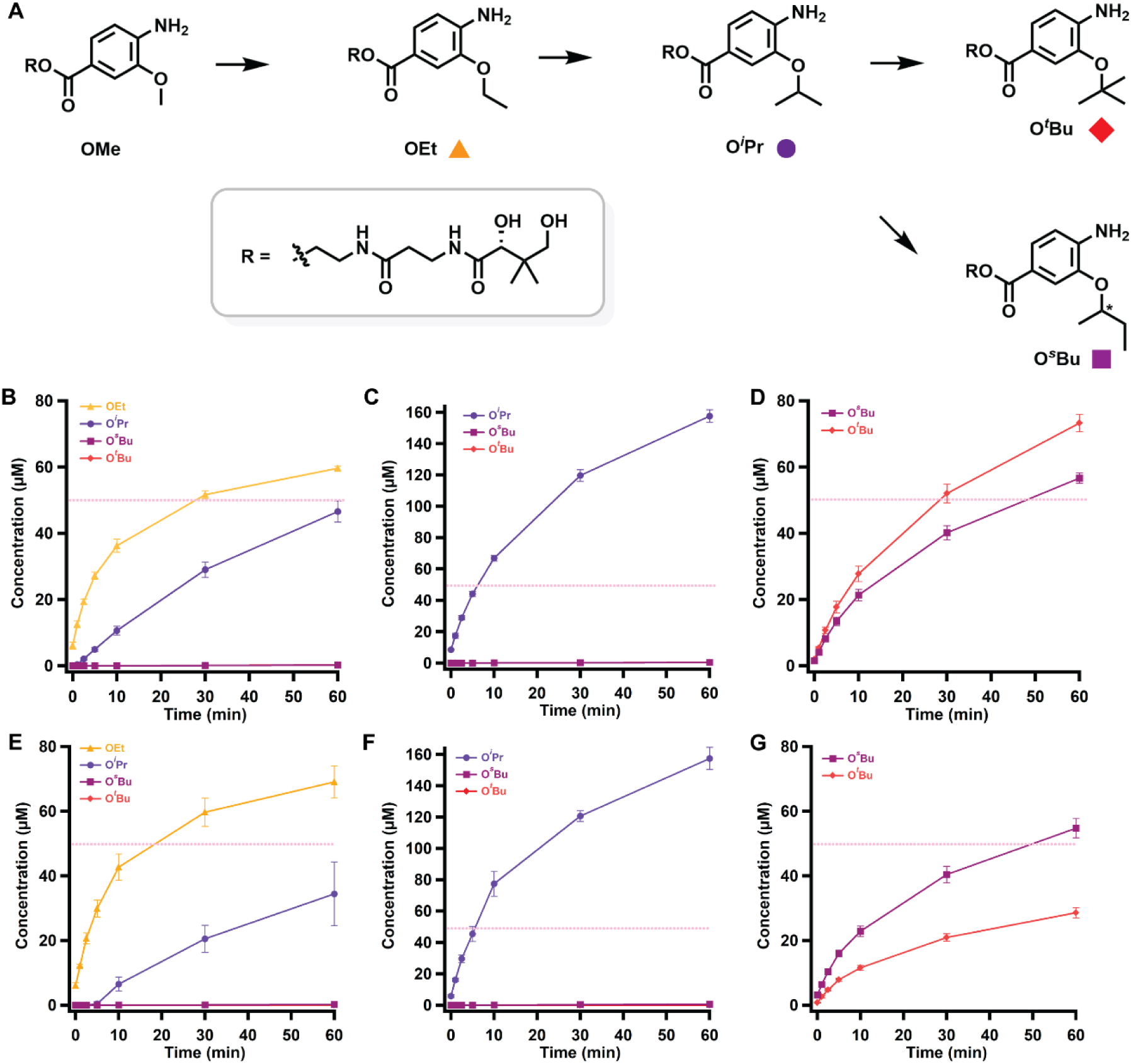
**A.** A scheme of the methylations performed by *cc*CysS and *cb*CysS. The colored squares under each structure represent the corresponding species in the graphs below. Quantification of methylated products in the *cb*CysS reactions using OMe-(**B**), OEt-(**C**), or O*^i^*Pr-containing species (**D**) as substrates; Quantification of methylated products in the *cc*CysS reactions using OMe-(**E**), OEt-(**F**), or O*^i^*Pr-containing species (**G**) as substrates. The dashed pink lines in each figure indicate the enzyme concentration. The asterisk in O*^s^*Bu indicates that the stereochemistry was not determined. The reactions contained 50 µM CysS, 500 µM SAM, 500 µM substrate, 2 mM Ti(III) citrate as a reductant, and L-tryptophan as an internal standard. The reactions were performed in triplicate. Error bars represent one standard deviation from the mean.

### Overall structure of *cc*CysS

*cc*CysS was crystallized under anoxic conditions in the presence of 5’-deoxyadenosine (5′dA), Met, and either the OMe, OEt, or O*^i^*Pr substrate, and the structures were determined to resolutions of 1.75 Å, 2.00 Å, and 1.95 Å, respectively (**Table S1**). Structures with the OMe or OEt substrates have well-defined substrate electron densities. However, the protein crystallized with the O*^i^*Pr substrate shows no electron density corresponding to the substrate. The overall architectures of all three structures are similar, composed of three domains (**Figure 3A**, topology diagram in **Figure S4**): a Cbl-binding domain (residues 1-185), a RS domain (residues 186-432), and a C-terminal domain (residues 433-630). We observe clear Fo-Fc electron density in the active site for the OMe and OEt substrates. The substrates are positioned between the 5′-carbon of 5′dA and the upper (distal) axial ligand of Cbl, suitable for H• abstraction and the subsequent radical attack on MeCbl (**Figure 3B**). While the upper ligand of Cbl is a water molecule, a Trp residue (Trp75) occupies the cofactor’s lower (proximal) axial face, residing 3.8 Å from the metal ion. This bulky residue, also observed at the same position in TokK (Trp76), creates a hydrophobic environment for Cbl’s cobalt ion, preventing water from approaching it and maintaining it in a pentacoordinated state (27). A similar arrangement was observed in Mmp10 but with a leucine residue in the lower axial position (31).

**Figure 3.**
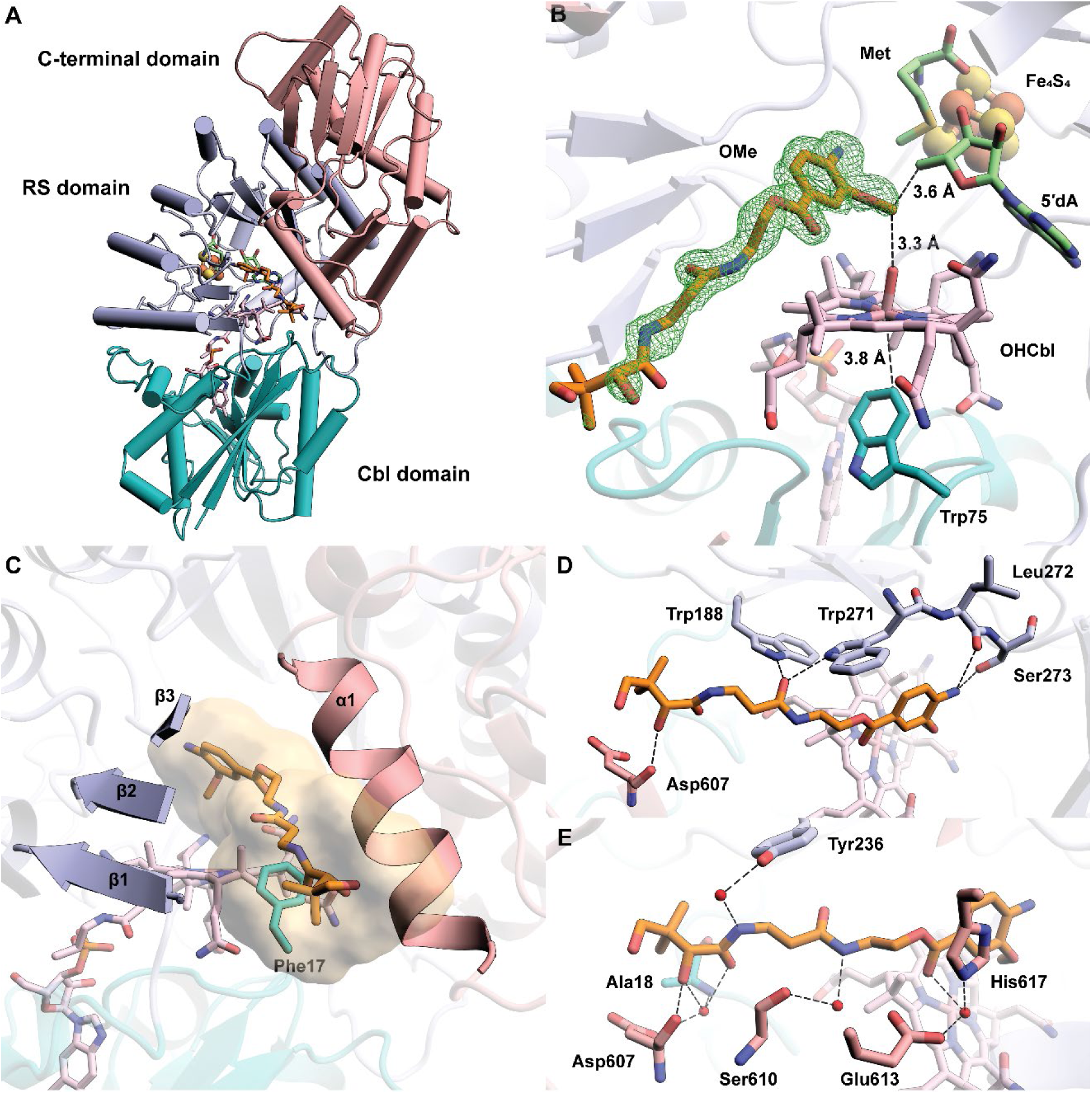
**A.** The overall structure of *cc*CysS illustrated as a ribbon diagram and colored by domains. The Cbl-binding domain is shown in teal. The RS domain is shown in light blue. The C-terminal domain is shown in pink; **B.** Closeup of Met, 5′dA, FeS cluster, Cbl, OMe substrate, and Trp75 within the active site of *cc*CysS. A simulated annealing composite omit 2Fo–Fc map (green mesh, contoured at 1.0 σ) is shown to provide an unbiased representation of the electron density for the OMe substrate. Only the major conformer of the substrate is displayed for clarity.Trp75 is the hydrophobic residue below the cobalt ion of Cbl. The distance between the cobalt ion and the closest atom of Trp75 is 3.8 Å; **C**. The substrate cavity is formed by the three β-strands of the RS domain, an α-helix of the C-terminal domain, and a hydrophobic residue (Phe17) from the Cbl-binding domain. The cavity is shown as a surface colored yellow. The cavity was generated by kvfinder (https://kvfinder-web.cnpem.br/) around the OMe substrate. The FeS cluster, Met, and 5′dA are not shown for clarity; **D.** Direct polar interactions between *cc*CysS and the OMe substrate; **E.** Through-water interactions between *cc*CysS and the OMe substrate.

In the *cc*CysS structure, the OMe substrate occupies a narrow channel leading from the protein’s surface into its active site. The channel is formed by the first three β-strands (β_1_-β_3_) from the TIM barrel of the RS domain, an α-helix (α_1_) from the C-terminal domain, and a hydrophobic residue (Phe17) from the Cbl-binding domain (**Figure 3C**). The β-strands from the RS domain contribute the most significant polar contacts with the substrate (**Figure 3D**). For example, the carbonyl group of Leu272 and the hydroxyl group of Ser273 form H-bonds with the aromatic amino group of the substrate, and the indole nitrogens of Trp188 and Trp271 interact with the same substrate carbonyl group. The α-helix (α1) from the C-terminal domain, which participates in forming the substrate cavity, has mostly through-water interactions with the substrate (**Figure 3E**). Asp607, located on α1, makes direct and through-water interactions with the secondary hydroxyl group in the pantetheine tail, highlighting the importance of this helix in substrate recognition. The OEt substrate has the same direct polar interactions with *cc*CysS as the OMe substrate (**Figure S5**).

### The OMe and OEt substrates bind similarly in the *cc*CysS active site

Clear electron density is observed for the OMe and OEt substrates in our structures. The electron density for the OMe substrate suggests two distinct conformations in a ratio of 2:1 (**Figure 4A, Figure S6**). The distance from C5′ of 5′dA to the carbon atoms of the methoxy group of both conformers is 3.6 Å. The distances from the carbon atoms of methoxy groups to the OH group of OHCbl are slightly different: 3.3 Å for the major conformer at an angle of 100.5° relative to C5 and 3.1 Å for the minor conformer at an angle of 103.0° relative to C5′. Both conformers seem plausible for H• abstraction and subsequent methylation. There is only one conformation of the OEt substrate, with distances of 3.9 Å and 3.1 Å from the methylene carbon of the ethoxy group to C5′ and the OH group of OHCbl, respectively (**Figure 4B**). The C5′-methylene-OHCbl angle is 99.3°, comparable with both angles observed in the two conformers of the OMe substrate. However, when overlaying the OMe structure with the OEt structure, the carbon of the methoxy group in the major OMe conformer occupies the equivalent position of the methylene carbon of the ethoxy group in the structure of *cc*CysS with the OEt substrate (**Figure S7A, B**). Likewise, overlaying the OMe-bound *cc*CysS structure with the substrate-bound TokK structure shows that the methoxy carbon in the major OMe conformer occupies the same spatial position as the methylene carbon to be methylated in the substrate of TokK (**Figure S7C, D**). An inspection of the environment around the OMe substrate identifies two bulky residues (Trp206 and Phe238) on the minor conformer side and one residue (Phe485) on the major conformer side (**Figure 4C**). These residues are in the same positions in the OEt structure. The greater steric hindrance from Trp206 and Phe238 prevents the ethoxy group from binding to that side. Thus, it is likely that the major conformation of the OMe substrate corresponds to the conformation for the first methylation, and all subsequent methylations occur on the side where Phe485 resides. Unfortunately, we were unable to determine the structure of *cc*CysS with the bound O*^i^*Pr substrate, which would have allowed us to visualize how the third methylation occurs and what governs the partitioning between the O*^s^*Bu and O*^t^*Bu products. The O*^s^*Bu product is formed by methylating the methyl carbon of the O*^i^*Pr substrate, while the O*^t^*Bu product is formed by methylating the methine carbon of the O*^i^*Pr substrate. We observe that during the reaction, the ratio of the two products remains constant, suggesting that at least two conformations of the O*^i^*Pr substrate exist.

**Figure 4.**
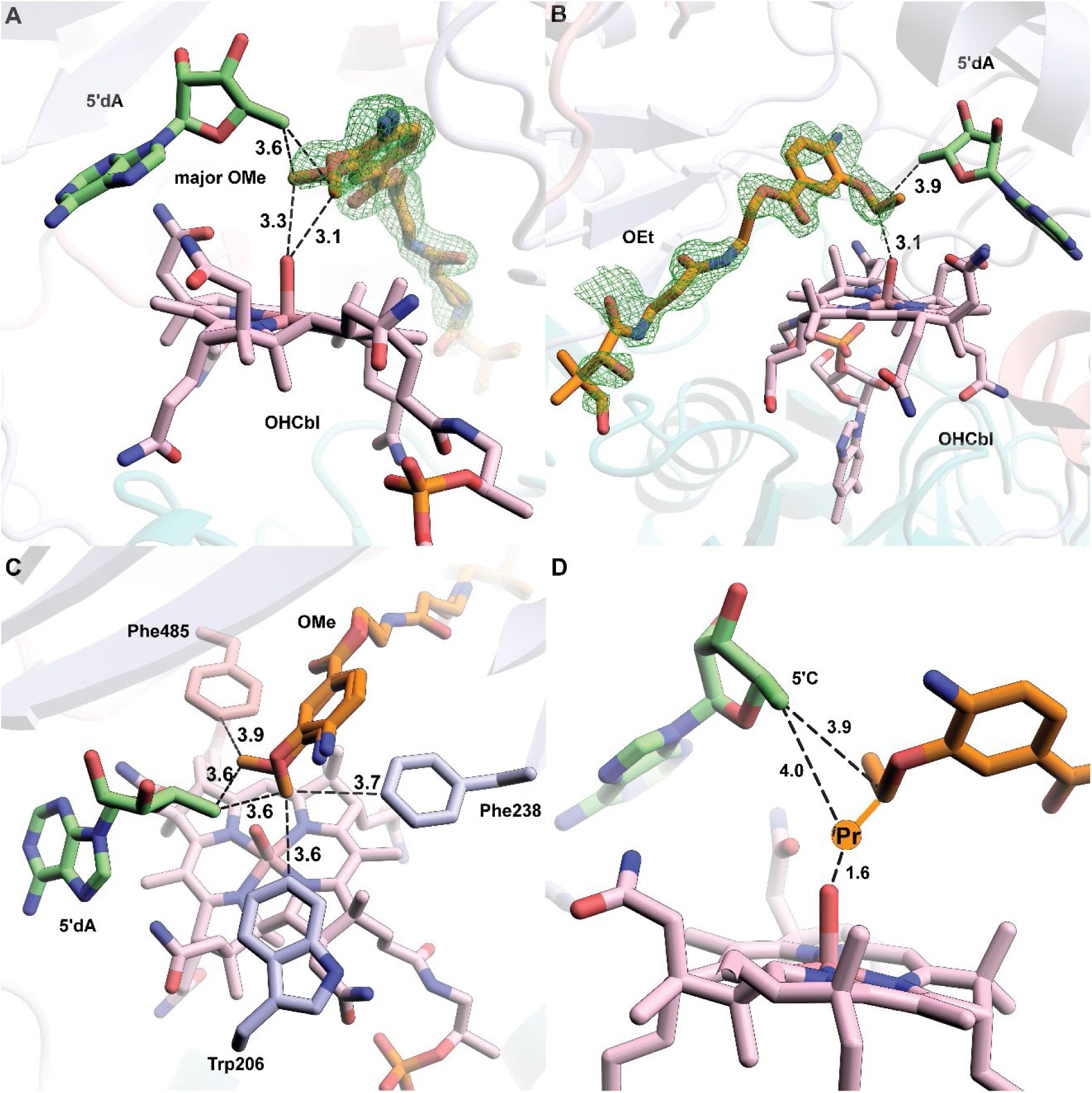
**A.** A simulated annealing composite omit 2Fo–Fc map (green mesh, contoured at 1.0 σ) is shown to provide an unbiased representation of the electron density for the OMe substrate. Both conformers of the OMe substrate are shown. **B.** A simulated annealing composite omit 2Fo–Fc map (green mesh, contoured at 1.0 σ) is shown to provide an unbiased representation of the electron density for the OEt substrate.; **C.** Trp206 and Phe238 provide steric hindrance for the minor conformer of OMe. Phe485 allows more space for the major conformer of the OMe substrate; **D.** Projection of the additional methyl group added to the methylene carbon in the OEt substrate and the respective distances from C5′ and the hydroxyl moiety of OHCbl. Small molecules, including substrates (OMe and OEt), cofactor (Cbl), and coproducts (Met and 5′dA), are shown in stick format and colored by atom type. Distances are indicated by dashed black lines and given in units of Å.

Phe485, which occupies different positions in the no-substrate structure versus the structures with substrates, appears to play a role in product partitioning. To study its function, Phe485 was replaced with either less bulky (Ala or Leu) or bulkier (Tyr or Trp) residues, and the variants were analyzed using the O*^i^*Pr substrate. The F485A variant produces no soluble protein, while the other three variants are isolated in moderate to good yields. As shown in **Figure S8**, F485Y and F485W yield fewer overall methylated products. The bulkier side chains of Tyr or Trp may sterically hamper the binding of the O*^i^*Pr substrate. The F485L variant, less bulky than wild-type (WT) *cc*CysS, produces methylated products in roughly equal amounts compared to the WT protein. Presumably, the additional space created by the F485L variant does not enhance the reaction. However, the ratio of the O*^s^*Bu and O*^t^*Bu products changes dramatically. WT *cc*CysS gives a 2:1 ratio of the *^s^*Bu/*^t^*Bu products. The ratio is slightly increased to 3:1 for the F485L variant. The F485Y variant yields a ratio of 5:1, while the F485W variant yields a ratio of 3:1. The change in product partitioning suggests that the conformational partitioning of the bound O*^i^*Pr substrate differs in each variant, presumably reflecting the bulkiness of the side chain of residue 485. However, like in WT *cc*CysS, all these variants still favor the O*^s^*Bu product over the O*^t^*Bu product.

### Trp271 is a gate for “ping” and “pong” steps

Cbl is bifunctional in Cbl-RSMs, participating in the S_N_2-dependent methylation of cob(I)alamin by SAM and the radical-dependent methylation of the substrate. Current structures of Cbl-RSMs cannot account for both types of chemistries occurring in a single active site without invoking significant conformational changes. Our *cc*CysS structure without substrate reveals a potential mechanism for how these enzymes regulate the two distinct chemistries. Compared with the OMe structure, the backbone of Trp271 in the no-substrate structure is unchanged, but the side chain rotates about 90° along the Cβ-Cγ bond. In a superposition of the two structures, this rotation results in a clash with the OMe substrate (**Figure 5A**). In the no-substrate structure, the entire Phe485 residue moves toward the entrance to the active site, where it would approach the OMe substrate in that structure but not clash with it. The movement of Phe485 is part of a larger conformational change involving a loop in the C-terminal domain (residues 480-485) (**Figure 5B**). Trp271, potentially in concert with Phe485, closes the substrate channel (**Figure 5C** vs. **Figure 3C, Figure S9**). The two orientations of Trp271 in the OMe and no-substrate structures indicate that it plays a dual role. It forms an H-bond with the substrate during radical-dependent methylation but blocks the substrate from binding to allow cob(I)alamin methylation to generate MeCbl. When Trp271 is changed to Ala, the variant shows much lower activity across all three methylations (**Figure S10**). W271H and W271Q, which can provide polar interactions similar to those of Trp, rescue activity slightly, supporting the role of Trp271 in substrate binding.

**Figure 5.**
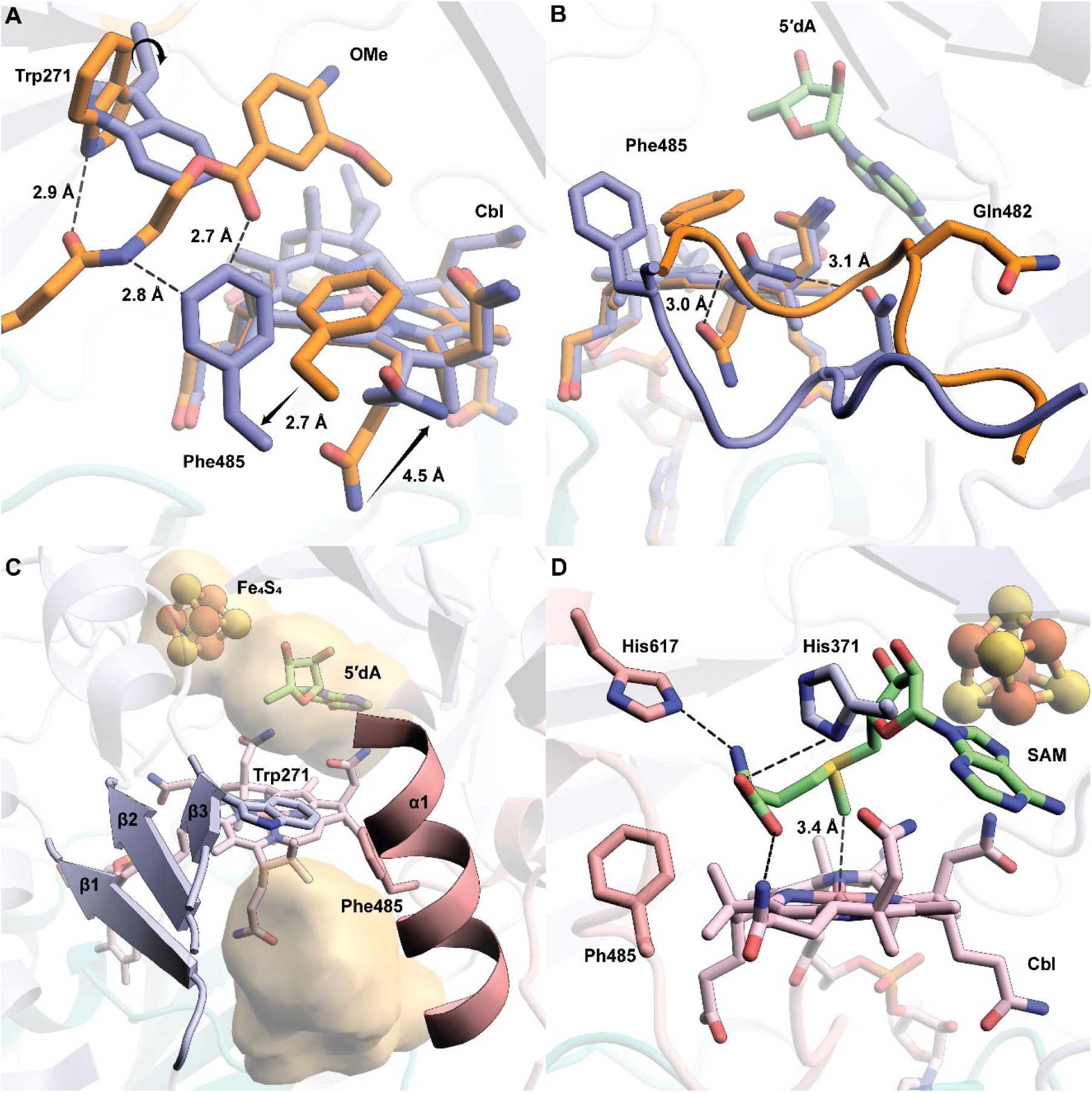
**A.** Overlay of the OMe structure with the no-substrate structure reveals the movement of Trp271, Phe485, and an acetamide side chain of Cbl. The black arrows indicate the movement of the indicated residues; **B.** The movements of Phe485 and the acetamide side chain of Cbl are due to the conformational change of a loop in the C-terminal domain of *cc*CysS. The residues, loops, and Cbl are colored orange for the OMe structure or blue for the no-substrate structure; **C.** The movement of Trp271 and Phe485 blocks the substrate channel created by β_1_-β_3_ from the RS domain and α_1_ from the C-terminal domain. The cavity is shown as a yellow surface. The cavity was generated by kvfinder (https://kvfinder-web.cnpem.br/); **D.** Docking model of SAM in the active site of the no-substrate structure shows the potential interaction between the methionine moiety of SAM and several residues from *cc*CysS and the acetamide side chain of Cbl. His617 and His371 are two potential residues that interact with the Met moiety of docked SAM. Both residues are colored by the domains they are from: His617 is from the C-terminal domain and colored pink; His371 is from the RS domain and colored blue.

In the absence of bound substrate, *cc*CysS should be primed to perform the S_N_2 methylation of Cbl. However, our current no-substrate structure has Met and 5′dA in the active site, which are not the products of Cbl methylation. To strengthen our hypothesis, a SAM molecule was docked into the active sites of both the OMe and no-substrate structures. During docking, we restricted the movement of the residues around the active site to verify that the active site can accommodate an appropriate pose of SAM for cob(I)alamin methylation. When docking SAM into the OMe structure with the substrate deleted, SAM shows only one conformation, in which it binds to the FeS cluster. This pose mimics Met (which binds to the FeS cluster) and 5′dA observed in the OMe structure, which reflects the reductive cleavage step (**Figure S11A**). When docking SAM into the no-substrate structure, the second lowest-energy conformation (the lowest is SAM bound to the FeS cluster) shows SAM extending its methionine moiety into the space created by the aforementioned loop movement (**Figure 5D**). In the docked model, His617, His371, and the acetamide side chain of Cbl make polar contacts with the methionine moiety of SAM. Notably, the methyl group of the docked SAM is right above the cobalt ion of Cbl, at a distance of 3.4 Å, representing an appropriate pose for cob(I)alamin methylation. When the OMe and no-substrate structures are overlayed, the methionine moiety clashes with Phe485 in the OMe structure (**Figure S11B**). These observations suggest that the movement of the Phe485-containing loop is cooperative with the rotation of Trp271. Trp271 closes the active site; however, the Phe485-containing loop moves to allow space for SAM to methylate cob(I)alamin.

## Discussion

In this study, we report three structures of *cc*CysS. One structure has no substrate, while the other two contain substrates for the first (OMe) and the second (OEt) methylations. The two substrates bind in almost identical poses in these two structures. Although we were unable to crystallize CysS with the O*^i^*Pr substrate, our biochemical studies identify a key residue (Phe485) that plays a role in the third methylation. When starting with the O*^i^*Pr substrate, changing Phe485 to other more or less bulky residues impacts the reaction rates or the product partitioning. The ratio of the two products from the third methylation by WT CysS is constant throughout time-dependent activity measurements. However, the ratio shifts when Phe485 is replaced with other residues. These data suggest that the O*^i^*Pr substrate binds in at least two distinct orientations, the ratio of which is governed to some extent by Phe485. The only substrate-bound structure of TokK contains the unmethylated compound. Assuming the same H-bonding networks are maintained as in our substrate-bound structures of *cc*CysS, the active site of TokK is large enough to allow three sequential methylations without a significant conformational change. A structural alignment identified a Tyr652 in the C-terminal domain of TokK, which is in a position corresponding to Phe485 in the C-terminal domain of *cc*CysS (**Figure S12**). However, no movement of the loop containing Tyr652 was observed in TokK, which contrasts with a significant loop movement observed in the structures of *cc*Cys with or without substrate (**Figure S13**). The third methylation by CysS may require a slight conformational change to accommodate the greater bulk, but this does not appear to be the case in TokK.

Trp271 provides one of the five direct H-bonds to the OMe or OEt substrates observed in the CysS structures, consistent with the diminished enzymatic activities of the W271A variant. Trp271 also appears to act as a gate. In the absence of substrate, Trp271 rotates its indole ring into the active site, preventing substrate binding. We suggest that this movement organizes the active site for the methylation of cob(I)alamin. This movement is accompanied by a conformational change of a C-terminal loop (residues 480-485), which creates a space into which the methionyl moiety of SAM can bind by interacting with His371, His617, and a propionamide appendage on the corrin ring of Cbl (**Figure S14**). Importantly, SAM cannot be docked into the same space in the CysS structure with substrate bound, even if the substrate is removed, highlighting the importance of the conformational change. Therefore, the X-ray crystal structure and docking experiments provide insight into how CysS allows for two different orientations of SAM to facilitate two distinct types of chemistry within one active site. As illustrated in **Figure 6**, for the S_N_2-dependent methylation of cob(I)alamin, Trp271 closes the active site to prevent the substrate from binding, while a loop simultaneously moves to provide space for SAM to bind in an appropriate conformation to methylate cob(I)alamin. Upon release of SAH, the indole ring of Trp271 rotates into a different conformation, opening the active site for the binding of another SAM molecule and the substrate. In this instance, the loop moves back to direct SAM to bind the FeS cluster for the radical-dependent methylation of the substrate. Upon release of the methylated product, the active site will be ready for the next round of catalysis, starting from Cbl methylation with a closed active site governed by Trp271 and Phe485.

**Figure 6.**
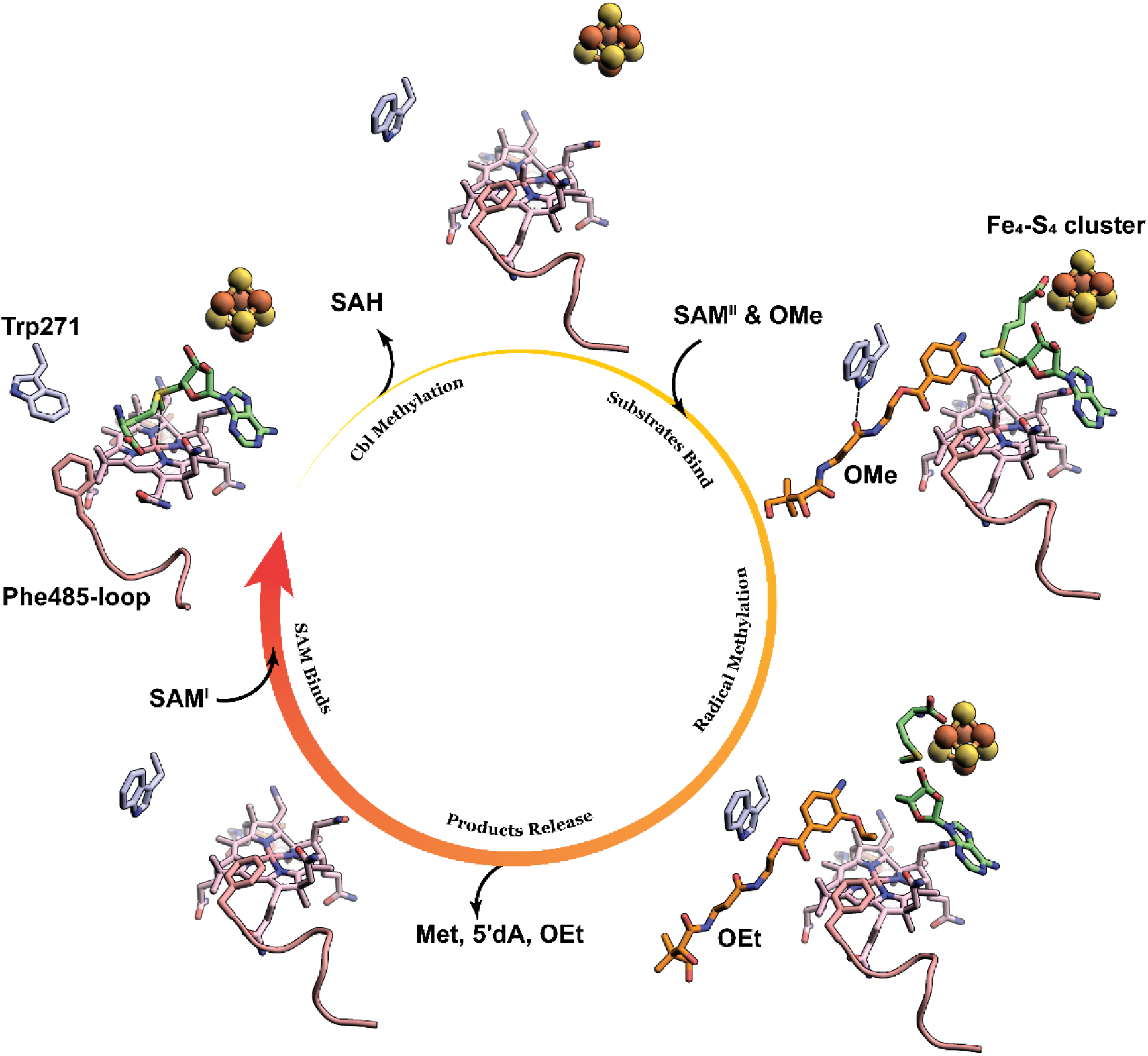
Illustration of the conformational changes of Trp271 and the Phe485-containing loop during the catalytic cycle. Trp271 is shown as light blue sticks. The Phe485-containing loop is shown as a pink cartoon. SAM, Met, and 5′dA are shown as green sticks. The OMe substrate and the OEt product are shown as orange sticks.

## Materials and Methods

All commercial materials were used as received unless otherwise noted. SAM was synthesized and purified as described previously (38). Perfluoropolyether cryo oil and crystal screening conditions were purchased from Hampton Research (Aliso Viejo, CA). Malonic acid, imidazole, boric acid (MIB) buffer were from Jena Bioscience (Jena, Germany). UV-visible spectra were recorded on a Varian Cary 50 spectrometer (Agilent Technologies, Santa Clara, CA) using the WinUV software package to control the instrument. High-resolution mass spectrometry (HRMS) was conducted on a Thermo Scientific Q Exactive HF-X hybrid quadrupole-Orbitrap mass spectrometer in line with a Vanquish Ultra-High Performance Liquid Chromatography (UHPLC). Data were collected and processed using Thermo Scientific Xcalibur 4.2.47 software.

### Overexpression and purification of CysS and its variants

The genes for CysS proteins from *Cystobacter sp. Cbv34* (UniProt ID: A0A0H4NV78, *cb*CysS) and *Corallococcus sp. CA054B* (UniProt ID: A0A3A8HCN5, *cc*CysS) and associated variants were codon-optimized for expression in *E. coli* (sequence in supporting information) and synthesized by Gene Universal (Newark, DE). Each gene was subcloned into a pSUMO vector using *Nde*I and *Xho*I restriction sites. The resulting plasmid was used to transform *E. coli* BL21 (DE3) competent cells containing plasmid pDB1282 and plasmid pBAD42-BtuCEDFB. A 200 mL starter culture in Luria-Bertani (LB) broth containing 50 mg/L kanamycin, 100 mg/L ampicillin, and 50 mg/L spectinomycin was inoculated with a single colony and shaken at 37 °C and 250 rpm for 12 h. 20 mL of the starter culture was used to inoculate 4 L of M9-ethanolamine medium containing 50 mg/L kanamycin, 100 mg/L ampicillin, and 50 mg/L spectinomycin and incubated at 37 °C with shaking at 180 rpm(34). Expression of the genes encoded on plasmid pDB1282 was induced at an OD_600_ of 0.3 with arabinose (8.0 g, 0.2% w/v). At an OD_600_ of 0.6, the flasks were placed in an ice-water bath for 0.5 h. Once cooled, IPTG was added to a final concentration of 0.5 mM, and iron chloride (FeCl_3_) was added to a final concentration of 25 µM. The cultures were incubated overnight for 18 h at 18 °C with shaking at 150 rpm. The cells were harvested, flash-frozen in liquid nitrogen, and stored at −80 °C until use.

45 g of frozen cell paste was resuspended in 150 mL of lysis buffer (50 mM HEPES, pH 7.5, 300 mM KCl, 10% (v/v) glycerol, 2.5 mM imidazole, and 10 mM β-mercaptoethanol (BME)) containing lysozyme (1 mg/mL), DNaseI (0.1 mg/mL), and an EDTA-free protease inhibitor tablet. For metallocofactor reconstitution during lysis, the resuspension mixture was supplemented with hydroxocobalamin acetate (1 mg/g of cell paste), cysteine (1 mg/mL), FeCl_3_ (0.8 mg/mL), and pyridoxal 5’-phosphate (1 mg/g of cell paste). The mixture was stirred at room temperature for 30 min until homogeneous. The resuspended cells were cooled on ice and subjected to six 45-s sonic bursts (50% output) using a QSonica instrument in a Coy anaerobic chamber with 8 min intermittent pauses. The lysate was centrifuged for 1 h at 50,000×g at 4 °C. The resulting supernatant was loaded onto Ni-NTA resin equilibrated in the lysis buffer. The resin was washed with 200 mL of the wash buffer (50 mM HEPES, pH 7.5, 300 mM KCl, 10% (v/v) glycerol, 20 mM imidazole, 10 mM BME) prior to elution of SUMO-CysS containing proteins with elution buffer (50 mM HEPES, pH 7.5, 300 mM KCl, 250 mM imidazole, 10% (v/v) glycerol, and 10 mM BME). The pooled eluate was concentrated by ultrafiltration using an Amicon Ultra centrifugal filter unit with a 10 kDa molecular weight cutoff membrane into approximately 2.5 mL. The resulting protein solution was exchanged into 3.5 mL cleavage buffer (75 mM HEPES, pH 7.5, 300 mM KCl, 10 mM imidazole, 15% (v/v) glycerol, and 1.5 mM BME) using a PD-10 column (GE Healthcare). Ulp1 (0.5 mL, 8 mg/mL, 4 mg), cobalamin (3.0 mg), FeCl_3_ (50 µL, 85 mM solution in 50 mM HEPES pH 7.5), and Na_2_S (50 µL, 100 mM solution in water) were added to the buffer-exchanged SUMO-CysS-protein (total volume 4.0 mL), and the resulting solution was incubated on ice overnight. The cleavage reaction mixture was loaded on a Ni-NTA column equilibrated with cleavage buffer, and the flow-through was collected, pooled, and concentrated using an Amicon Ultra centrifugal filter unit with a 10 kDa molecular weight cutoff membrane. The resulting CysS proteins were further purified by size-exclusion chromatography on a HiPrep 16/60 S200 column housed in an anaerobic chamber, eluting isocratically in S200 buffer (50 mM HEPES, pH 7.5, 400 mM KCl, 10% glycerol, and 4 mM DTT). Fractions corresponding to monomeric CysS proteins were pooled, concentrated, frozen in liquid nitrogen, and stored in liquid nitrogen.

### Crystallization of *cc*CysS

All manipulations were carried out in a Coy (Grass Lake, MI) anaerobic chamber at room temperature. The protein solutions contained 10 mg/mL *cc*CysS, 10 mM HEPES, pH 7.5, 2 mM Met, 2 mM 5′dA, and 2 mM OMe, OEt, or O*^i^*Pr substrate. All crystals were large brown plates obtained within a week using the hanging-drop vapor-diffusion method. *cc*CysS + the OMe substrate crystals formed in solution A (100 mM MES buffer, pH 6.0, 200 mM MgCl_2_, 25% w/v PEG 6000) after mixing 2 µl of the protein solution with 2 µl of solution A in the hanging drop and equilibrating against 0.6 mL of solution A. Crystals were briefly dipped in cryoprotectant, mounted on nylon loops, and flash-cooled in liquid nitrogen. The cryoprotectant was made by mixing 100 µL of solution A, 100 µL of 50% (w/v) PEG 6000, and 8 µL of an aqueous solution of the OMe substrate (70 mM). *cc*CysS + the OEt substrate crystals formed in solution B (100 mM MES buffer, pH 6.0, 200 mM LiCl, 25% w/v PEG6000) after mixing 2 µl of the protein solution with 2 µl of solution B in the hanging drop and equilibrating against 0.6 mL of solution B. Crystals were briefly dipped in cryoprotectant, mounted on nylon loops, and flash-cooled in liquid nitrogen. The cryoprotectant was made by mixing 100 µL of solution B, 100 µL of 50% (w/v) PEG6000, and 10 µL of an aqueous solution of the OEt substrate (50 mM). *cc*CysS + the O*^i^*Pr substrate crystals formed in solution C (100 mM MES buffer, pH 6.0, 200 mM NaCl, 15% w/v PEG6000) after mixing 2 µl of the protein solution with 2 µl of solution C in the hanging drop and equilibrating against 0.6 mL of solution C. Crystals were briefly dipped in cryoprotectant, mounted on nylon loops, and flash-cooled in liquid nitrogen. The cryoprotectant was prepared by mixing 100 µL of solution C, 100 µL of 50% (w/v) PEG 6000, and 10 µL of an aqueous solution of the O*^i^*Pr substrate (50 mM).

### X-Ray data collection, processing, structure-determination, and refinement

X-ray diffraction datasets of *cc*CysS crystals were collected at λ = 0.97872 Å with 360° of data measured using a 1° oscillation at APS Beamline 21-ID-F (*cc*CysS+the OMe substrate); λ = 1.03317 Å with 360° of data measured using a 0.4° oscillation at APS Beamline 23-ID-B (*cc*CysS+the OEt substrate); λ = 1.03580 Å with 360° of data measured using a 0.25° oscillation at ALS Beamline 8.2.2 (*cc*CysS+the O*^i^*Pr-containing substrate). All datasets were processed using the HKL2000 package. CC1/2 (greater than 0.7) were primarily used to decide the resolution limits. Structures were determined by molecular replacement using the program PHASER (39–42). Model building and refinement were performed with Coot and phenix.refine (39, 43). Data collection and refinement statistics are shown in **Table S1**. Figures were prepared using PyMOL (44, 45). An Alpha fold model (https://alphafold.ebi.ac.uk/entry/A0A3A8HCN5) was used for molecular replacement using Phenix Phaser-MR (39). Iterative manual model building and refinement were performed in Coot and Phenix (39, 43). *R_free_* flags were determined by Phenix using the default settings (10% or up to 2000 reflections). Geometric restraints for 5′dA and cobalamin were obtained from the Grade Web Server (Global Phasing). Geometric restraints for the OMe and OEt substrates were obtained by eLBOW using the eLBOW AM1 QM method (46). In the later stages of refinement, Translation/Libration/Screw (TLS) parameters were additionally used. The TLS parameters were generated from TLSMD ( http://skuld.bmsc.washington.edu/~tlsmd/index.html). We selected five TLS groups based on the TLSMD score versus group-number analysis, where the five-group partition represented the point of maximal improvement. These groups correspond to residues 5–181, 182–186, 187–421, 422–515, and 516–704. Restraints for [Fe_4_S_4_] clusters were based on the structure of *M. thermoacetica* carbon monoxide dehydrogenase/acetyl-CoA synthase (PDB ID: 3I01) (47). For the OMe substrate, both conformers were given an arbitrary occupancy of 0.5 prior to refinement. The resulting occupancies of the major (0.57) and minor (0.2) conformers were calculated during the refinement process.

### Docking studies

Molecular docking was performed using Maestro 9.1 and the Glide module of the Schrödinger Suite (Schrödinger, LLC, New York, NY, USA). The initial structure employed in this study was generated from the X-ray crystal structure of *<u>cc</u>*CysS with no substrate bound (PDB ID: 9N1D). The protein preparation wizard was used to prepare the initial structure of *cc*CysS by adding hydrogen atoms, removing water, fixing the cofactors in the [Fe_4_S_4_]^+^ and cob(I)alamin states. A restrained minimization of the protein structure was performed using the OPLS_2005 force field to reposition side chains, hydroxyl groups, and resolve potential steric clashes with a predefined Root Mean Square Deviation (RMSD) tolerance of 0.3 Å. The receptor grid generation module was used to generate a 10 Å receptor grid using the prepared protein structure over the co-crystallized ligand molecules, as mentioned. To examine active site rearrangements and their effect on SAM’s conformation, the CysS structure with the OMe substrate bound (PDB ID: 9N1B) was also used for grid preparation, while the OMe substrate was removed from the active site during the initial preparation phase. For docking, we used ligands from either a crystal structure or a chemical database. We built all ligands using Maestro’s build panel and optimized them to lower-energy conformers with LigPrep. LigPrep also helps to assign chirality and convert compounds to 3D structures while generating low-energy conformers and relevant ionization/tautomeric states.

Glide docking of the proposed molecules was performed using the pre-prepared receptor grid and ligand structures. The Glide docking program evaluated the favorable interactions between the ligands and the receptor. All docking was performed in extra precision (XP) mode using the OPLS_2005 force field. To soften the interaction potential for the nonpolar part of the ligand, van der Waals radii were scaled to 0.8 with a partial charge cutoff of 0.15 for charged atoms. This docking process used a flexible docking method that automatically generated multiple conformations for each ligand while keeping the receptor rigid. The resulting ligand poses were evaluated through a series of hierarchical filters to analyze their interactions with the receptor. The algorithm identified favorable hydrophobic interactions, hydrogen bonds, and metal-ligation interactions, while assigning penalties for steric clashes. Poses that passed these initial filters were carried to the final stage, where the grid approximation of OPLS non-bonded ligand-receptor interaction energy was calculated and minimized. The minimized poses were then re-scored using the Glide Score scoring function for final evaluation. The reliability of the docking process was assessed by comparing how closely the lowest-energy pose (binding conformation) of the co-crystallized ligand, predicted by the Glide score (G Score), matched the experimental binding mode in the substrate-bound complex as determined by X-ray crystallography. The extra precision (XP) Glide docking method was validated by first removing the co-crystallized ligand from the protein’s binding site and then redocking it into the same site. The accuracy of the prediction was evaluated based on hydrogen-bonding interactions and the RMSD between the predicted conformation and the experimentally observed X-ray crystallographic structure.

### Enzymatic reactions

All activity assays were conducted in triplicate and contained 50 µM CysS, 500 µM 3-alkoxy-4-aminobenzoic acid-containing substrates, 500 µM SAM, 2 mM Ti(III) citrate, 250 mM KCl, and 90 µM tryptophan as an internal standard in 75 mM HEPES, pH 7.5. Reactions were quenched by adding a twofold volume of methanol at each time point. All reactions were monitored over 1 h at room temperature. The quenched samples were vortexed and centrifuged at 14,000 ×g and 4 °C for 45 min, and the resulting supernatants were analyzed by HRMS-UHPLC.

Quantification of all methylated products was conducted on an Agilent Technologies 1290 Infinity II series UHPLC system coupled to a 6470 QQQ Agilent Jet Stream electrospray-ionization mass spectrometer. Separation of the products was performed at 24.5 °C on an Agilent Zorbax Extend-C18 RRHD column (2.1 mm × 50 m, 1.8-µm particle size) equilibrated with 98% solvent A (0.1% formic acid, pH 2.5) and 2% solvent B (acetonitrile). One microliter of the quenched supernatant was injected onto the column, and analytes were separated using the following gradient: 2% solvent B from 0 to 2 min, 2% to 50% solvent B from 2 to 8 min, 50% to 60% solvent B from 8 to 11 min, 60% to 100% solvent B from 11 to 13 min, 100% solvent B from 13 min to 16 min, 100% to 2 % solvent B from 16 to 18 min, and 2% solvent B from 18 to 20 min. All analytes were detected in positive mode using multiple-reaction monitoring. A standard curve of all predicted methylation products (1 µM to 200 µM) containing 90 µM tryptophan (internal standard) was prepared for product quantification using the Agilent MassHunter Quantitative Analysis 10.1 software.

## Acknowledgments

We thank Dr. David F. Iwig for acquiring accurate mass data and Dante A. Serrano for optimizing the mass spectrometry method. This work was supported by NIH GM-122595 and the Eberly Family Distinguished Chair in Science (S.J.B.). S.J.B. is an investigator of the Howard Hughes Medical Institute.

This research used resources of the Advanced Photon Source, a DOE Office of Science User Facility operated for the DOE Office of Science by Argonne National Laboratory under contract no. DE-AC02-06CH11357. Use of GM/CA@APS has been funded in whole or in part with Federal funds from the National Cancer Institute (ACB-12002) and the National Institute of General Medical Sciences (AGM-12006). The Eiger 16M detector at GM/CA-XSD was funded by NIH grant S10 OD012289. Use of LS-CAT Sector 21 was supported by the Michigan Economic Development Corporation and the Michigan Technology Tri-Corridor (grant 085P1000817). This research also used the resources of the Berkeley Center for Structural Biology supported in part by the HHMI. The Advanced Light Source is a DOE Office of Science User Facility under contract no. DE-AC02-05CH11231. The ALS-ENABLE beamlines are supported in part by the NIH, National Institute of General Medical Sciences, grant P30 GM124169.

This article is subject to HHMI’s Open Access to Publications policy. HHMI lab heads have previously granted a non-exclusive CC BY 4.0 license to the public and a sublicensable license to HHMI in their research articles. Pursuant to those licenses, the author-accepted manuscript of this article can be made freely available under a CC BY 4.0 license immediately upon publication.

