## Supplemental Information for "Structural Basis for Iterative Methylation by a Cobalamin-dependent Radical S-Adenosylmethionine Enzyme in Cystobactamids Biosynthesis"

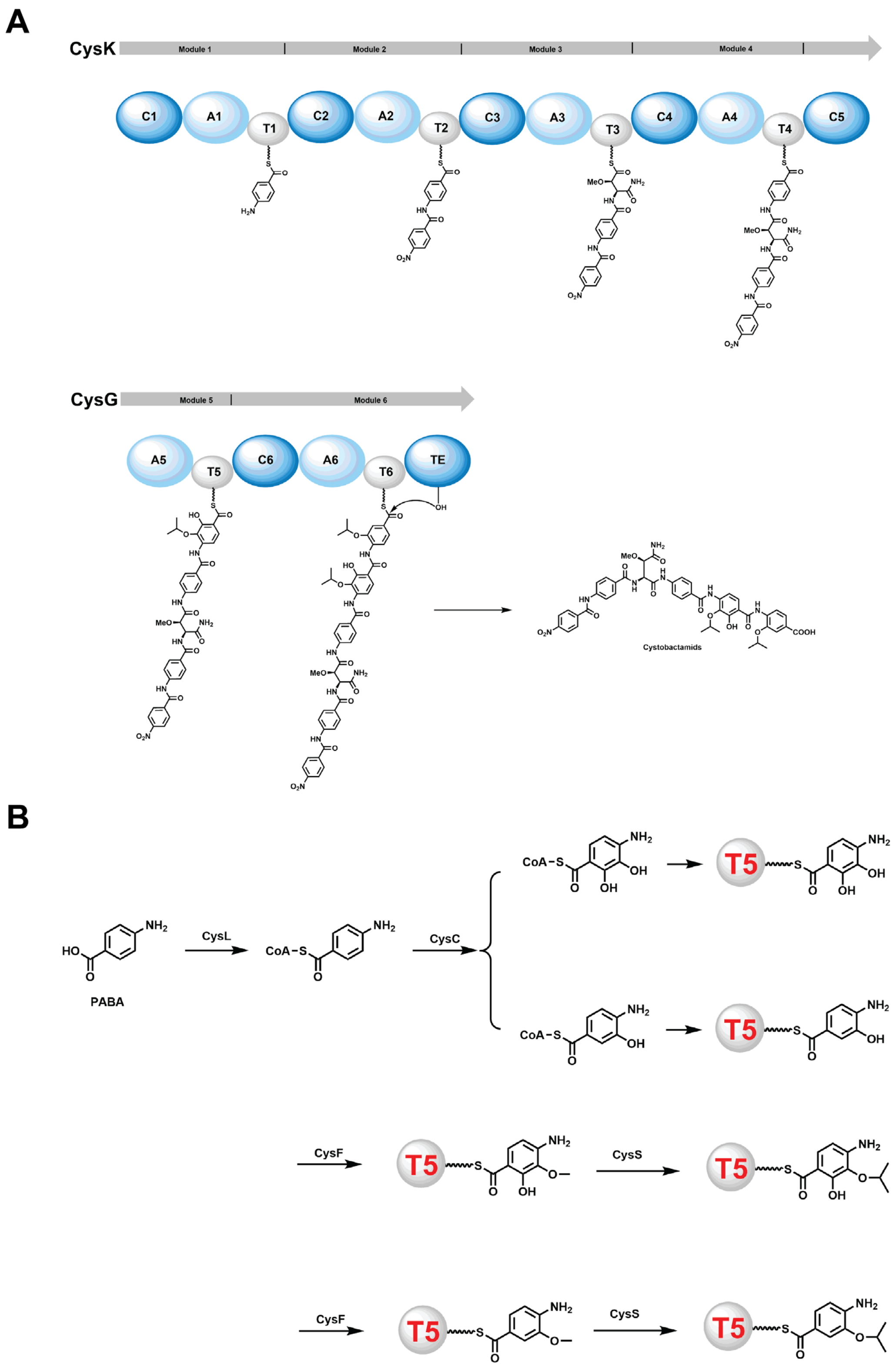


**Figure S1. A.** Proposed assembly of cystobactamids by two NRPS, CysK and CysG; **B.** Tailoring of PABA by CysC, CysF, and CysS before its incorporation into cystobactamids.

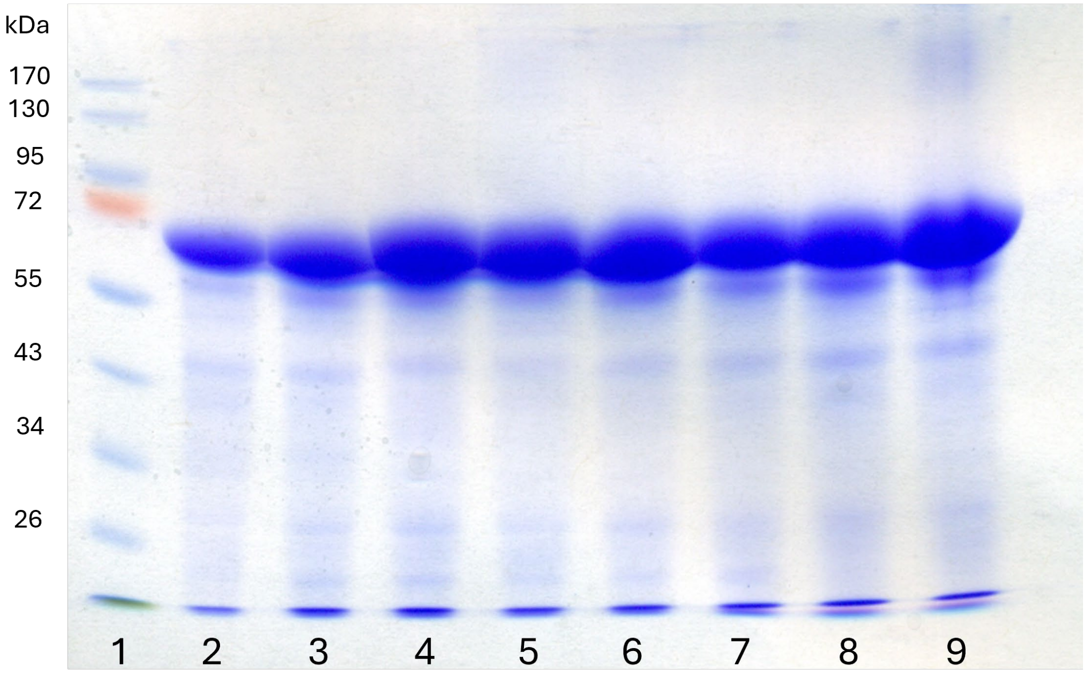


**Figure S2.** SDS-PAGE gel of purified CysS and its variants. Sample lanes: 1:ladder, 2:*cb*CysS, 3:*cc*CysS, 4:*cc*CysS-W271A, 5:*cc*CysS-W271H, 6:*cc*CysS-W271Q, 7:*cc*CysS-F485W, 8:*cc*CysS-F485Y, 9:*cc*CysS-F485L.

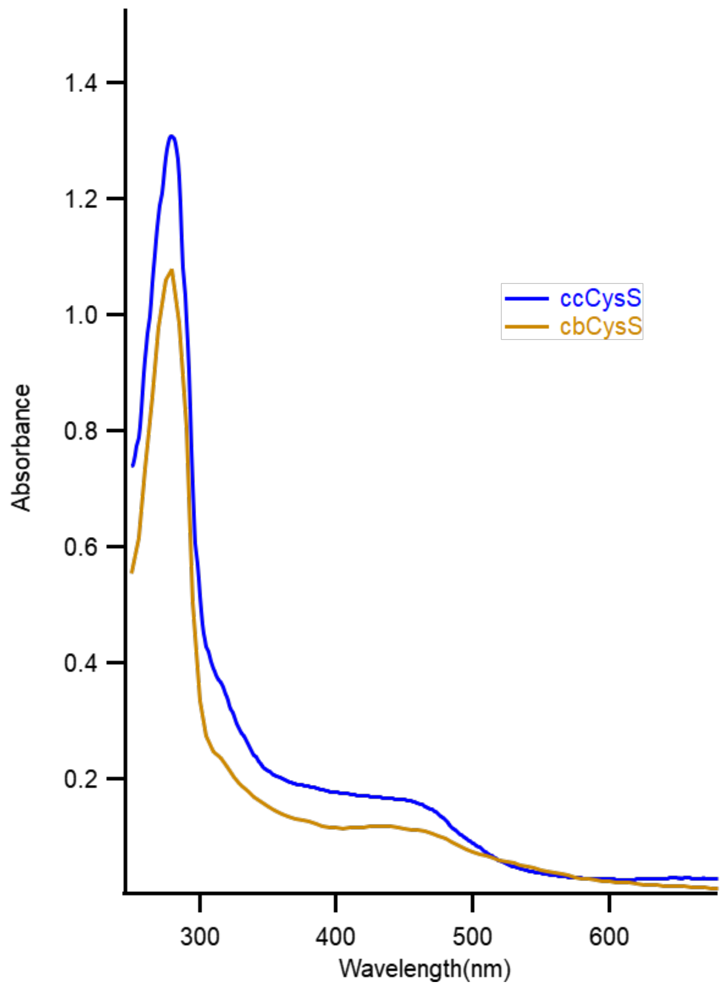


**Figure S3.** UV-vis spectra of *cb*CysS (17.65 μM) and *cc*CysS (17.34 μM).

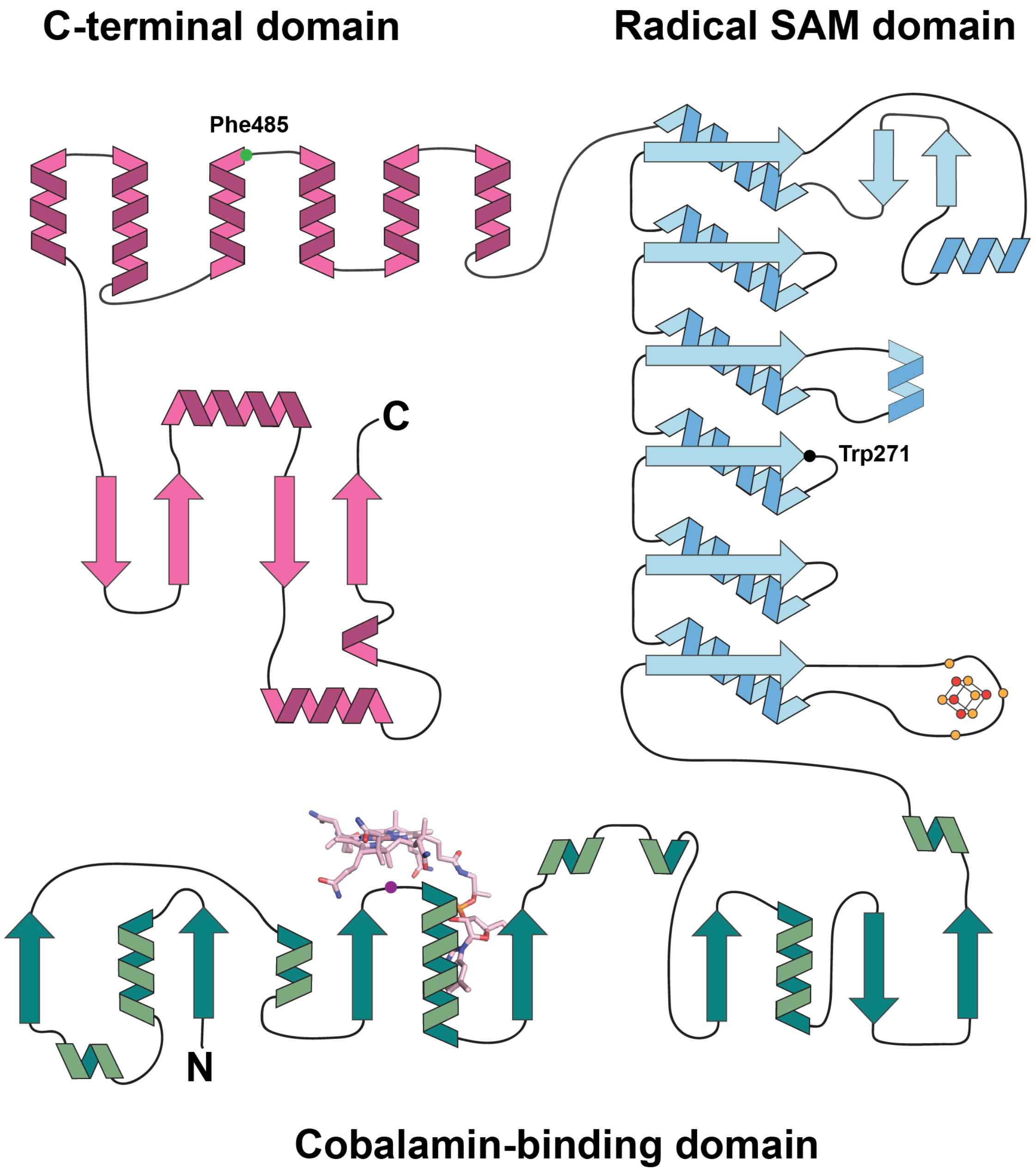


**Figure S4.** Topology diagram of *cc*CysS depicting domains. The cobalamin-binding domain is shown in teal. The RS domain is shown in light blue. The C-terminal domain is shown in pink. Three yellow dots surrounding the iron-sulfur cluster represent cysteine residues coordinating the cluster. Purple dot represents Trp75, the hydrophobic residue underneath the cobalt ion of cobalamin. Black dot represents Trp271. Green dot represents Phe485.

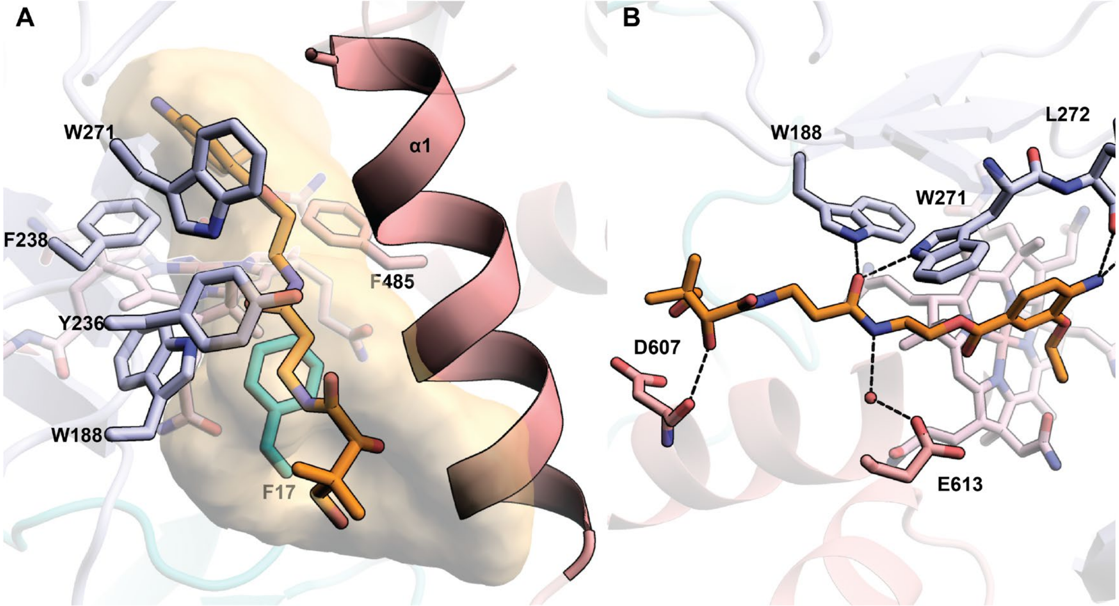


**Figure S5. A.** The substrate cavity in the *cc*CysS structure with the OEt substrate bound. The substrate cavity is formed by various hydrophobic residues from all three domains and α1 from the C-terminal domain. The hydrophobic residues are colored according to their domains. Trp188, Tyr236, Phe238, and Trp271 are from the RS domain and are colored light blue. Phe17 is from the Cbl-binding domain and is colored teal. Phe485 is from the C-terminal domain and is colored pink. The cavity is shown as a yellow surface. The cavity was generated by kvfinder (https://kvfinder-web.cnpem.br/) around the OEt substrate. **B.** Polar interactions between *cc*CysS and OEt substrate. OEt maintains all the key interactions observed in OMe-*cc*CysS structure.

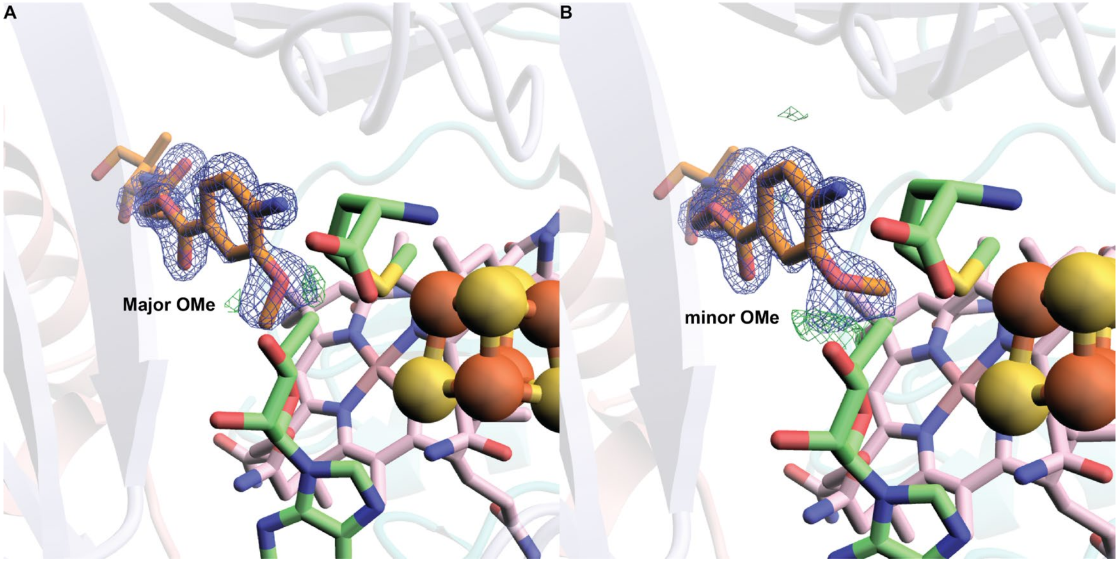


**Figure S6.** Individual electron density supporting alternate conformations of the OMe substrate in the OMe–*cc*CysS structure. Models were refined containing only the major (**A**) or only the minor (**B**) conformer of the substrate. The 2Fo–Fc electron density (blue mesh) is shown contoured at 1.5σ to illustrate the modeled conformer. The positive Fo–Fc difference electron density (green mesh, contoured at +3.0σ) reveals residual density corresponding to the alternate conformer in each case. To generate these maps, one conformer was omitted, and the model was refined to avoid model bias.

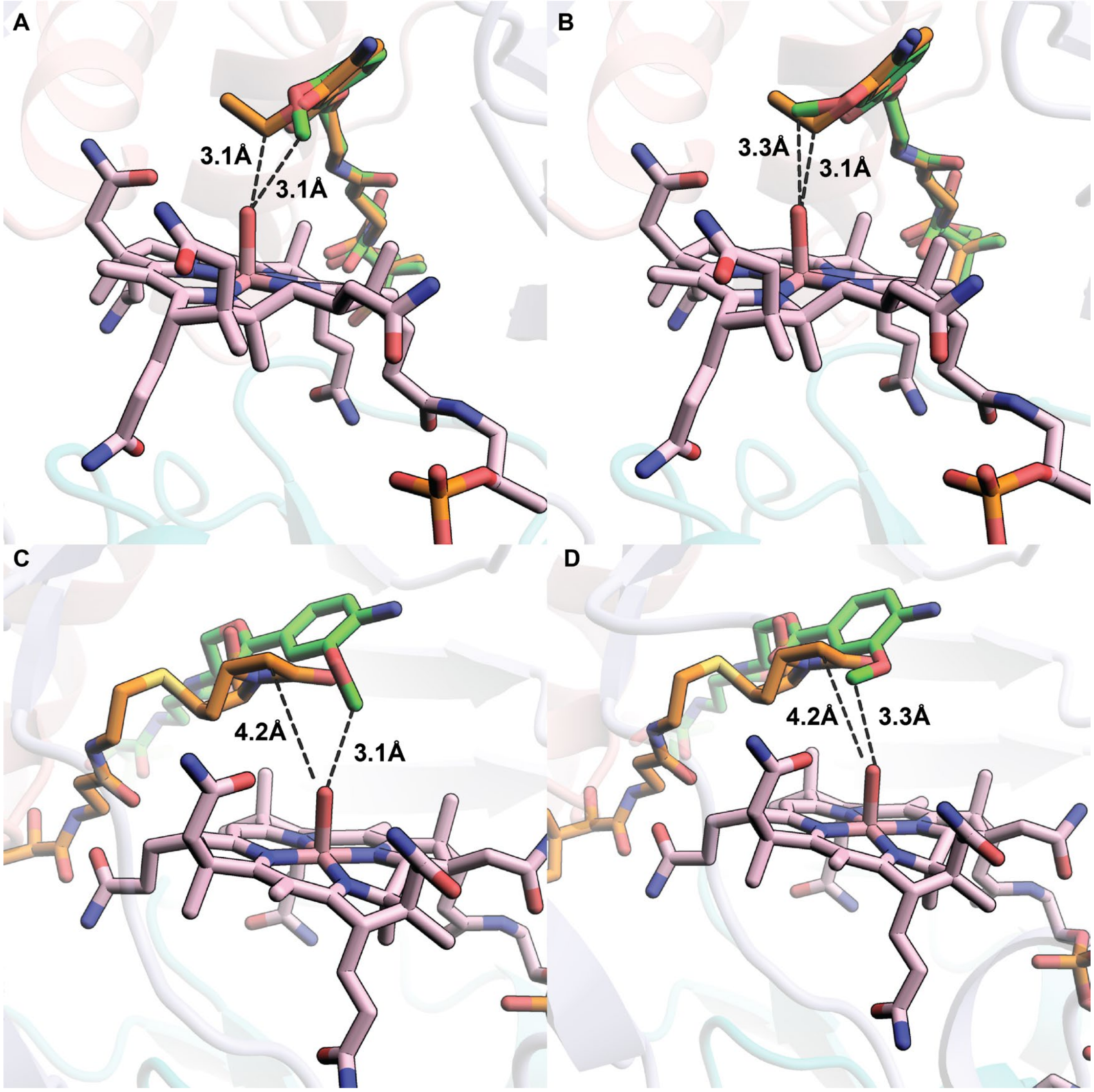


**Figure S7.** Overlay of the minor conformer of the OMe substrate with the OEt substrate (**A**) or the TokK (PDB ID: 7KDY) substrate (**C**). Overlay of the major conformer of the OMe substrate with the OEt substrate (**B**) or the TokK substrate (**D**).

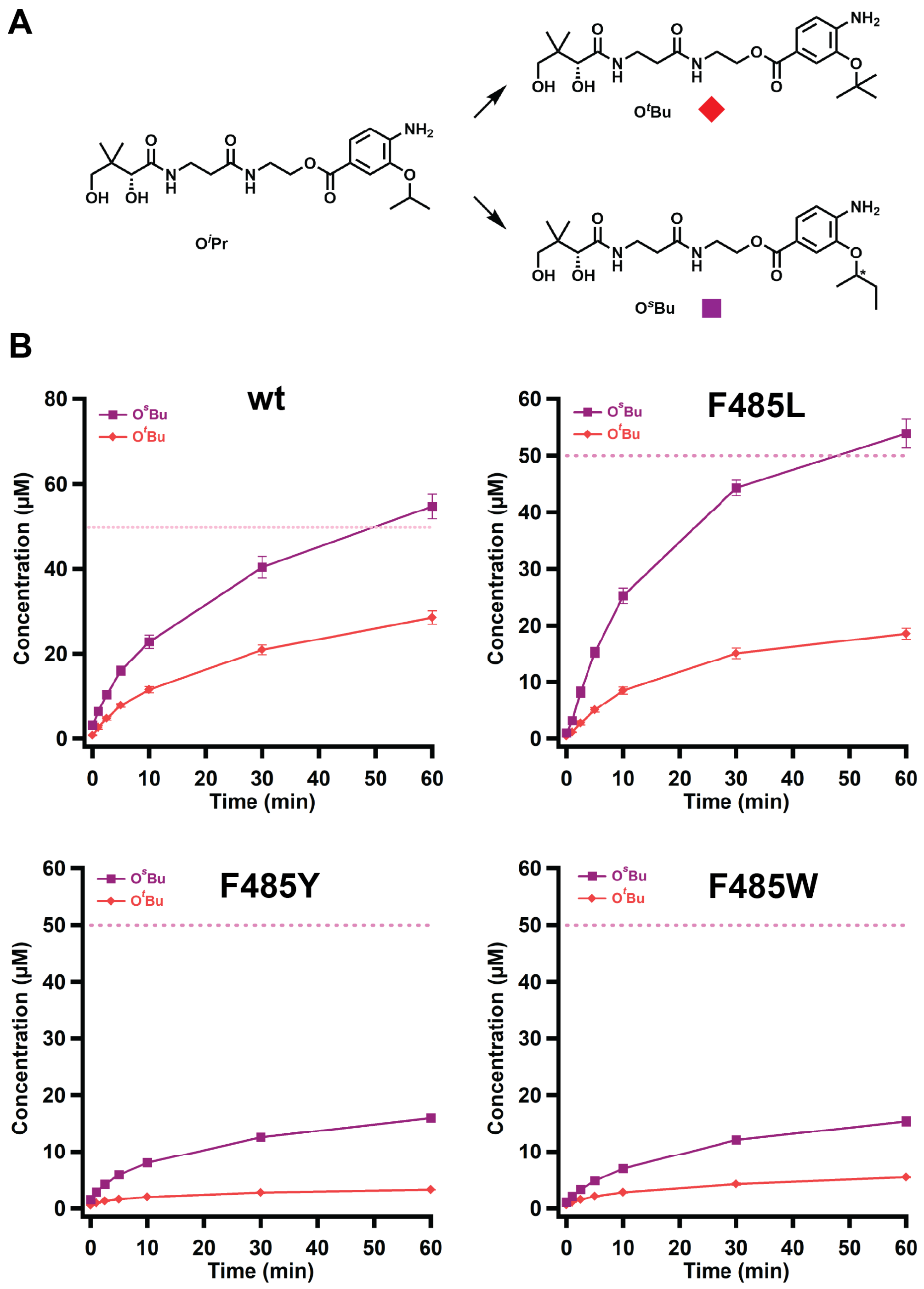


**Figure S8 A.** Scheme of the methylations performed on the O$^{i}$Pr substrate by *cc*CysS and its variants. The colored squares under each structure represent the corresponding species in the graphs below. **B.** Quantification of O$^{s}$Bu or O$^{t}$Bu products in the reactions with wt ccCysS, and the F485L, F485Y, and F485W variants. The dashed pink lines in each figure indicate the enzyme concentration. The asterisk in O$^{s}$Bu indicates that the stereochemistry was not determined. The reactions contained 50 μM CysS, 500 μM SAM, 500 μM substrate, 2 mM Ti(**III**) citrate as a reductant, and L-tryptophan as an internal standard. The reactions were performed in triplicate. Error bars represent one standard deviation from the mean.

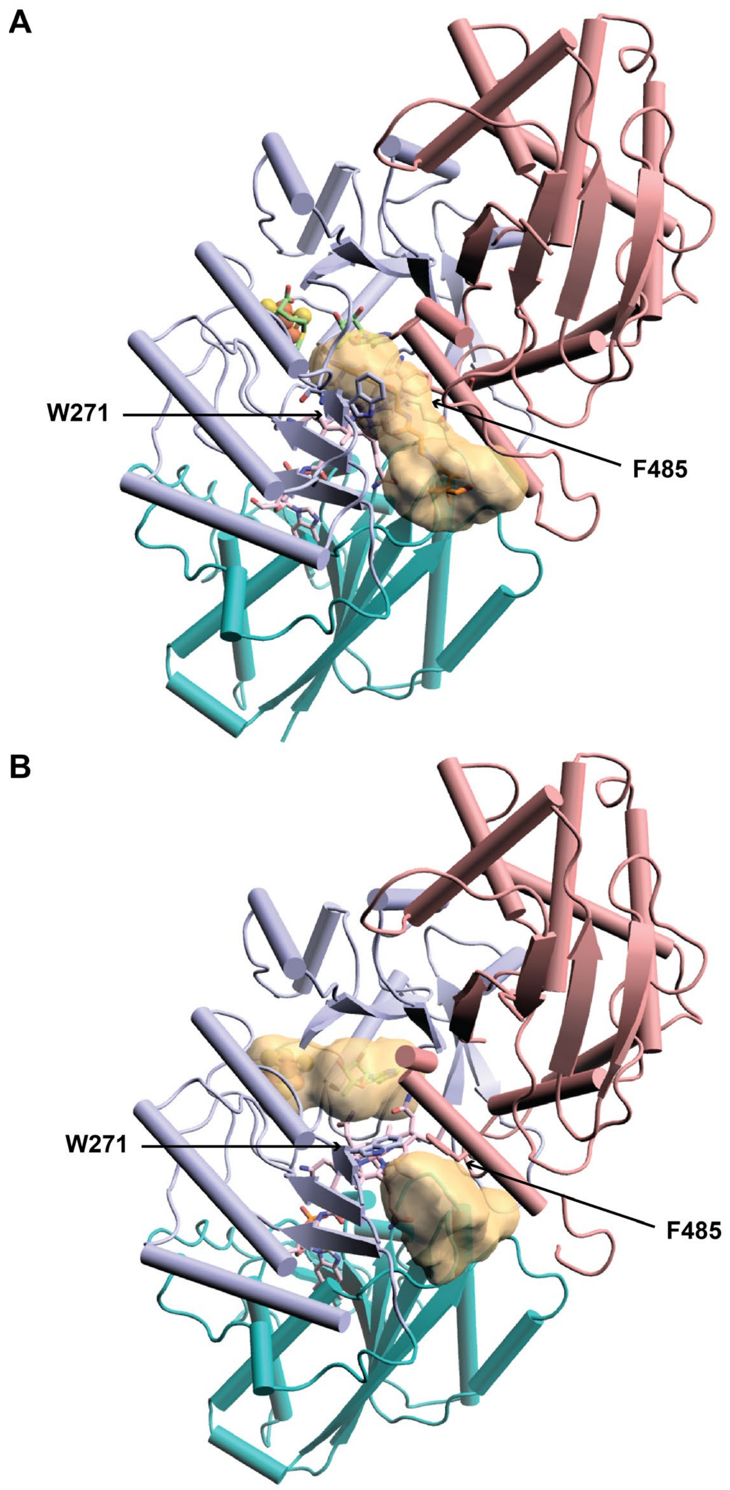


**Figure S9.** Comparison of the substrate cavities of OMe-*cc*CysS structure (**A**) and no-substrate-*cc*CysS structure (**B**). The cavity was generated by kvfinder (https://kvfinder-web.cnpem.br/). The movement of Trp271 and Phe485 closed the entrance of the cavity.

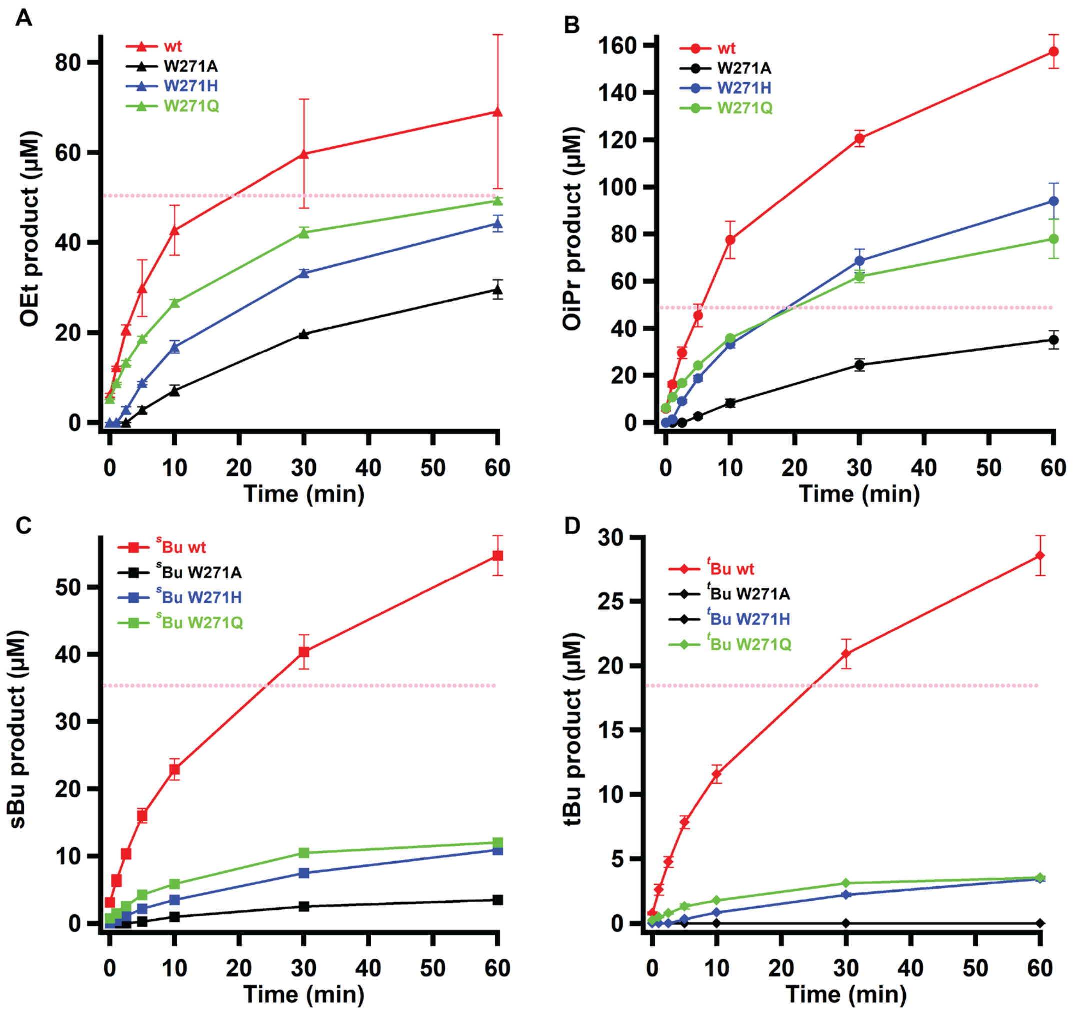


**Figure S10.** Quantification of the OEt product using the OMe substrate (**A**), the O$^i$Pr product using the OEt substrate (**B**), the O$^s$Bu (**C**) and O$^t$Bu products (**D**) using the O$^i$Pr substrate, in reactions with wt *cc*CysS, or the W271A, W271H, and W271Q variants. The dashed pink lines in each figure indicate the enzyme concentration. The reactions contained 50 μM CysS, 500 μM SAM, 500 μM substrate, 2 mM Ti(III) citrate as a reductant, and L-tryptophan as an internal standard. The reactions were performed in triplicate. Error bars represent one standard deviation from the mean.

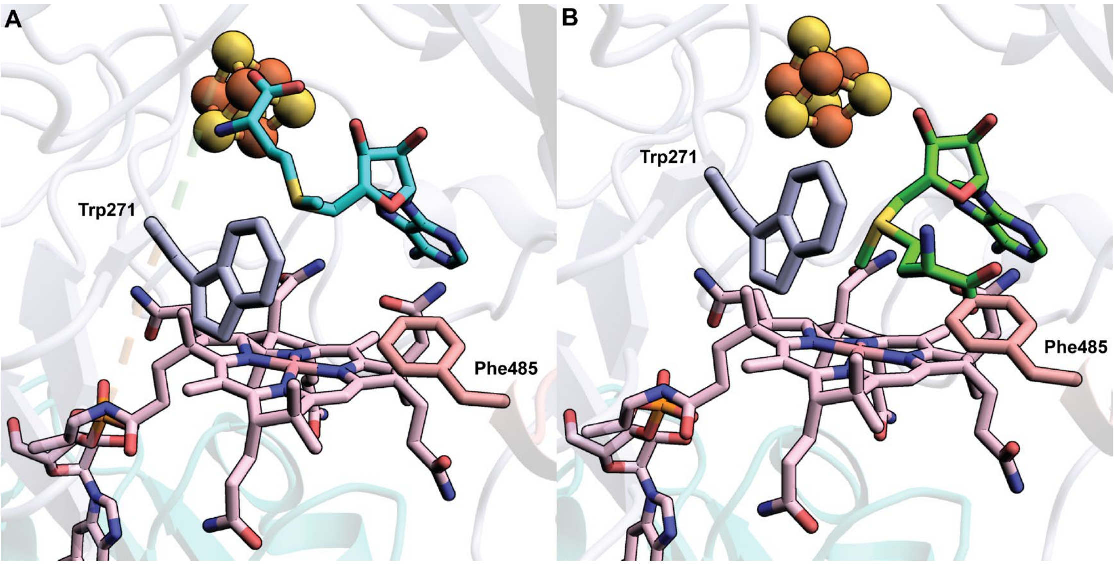


**Figure S11. A.** SAM docked into the OMe structure. The OMe substrate was deleted before docking; **B.** SAM docked into a no-substrate structure clashes with Phe485 if placed in the OMe structure.

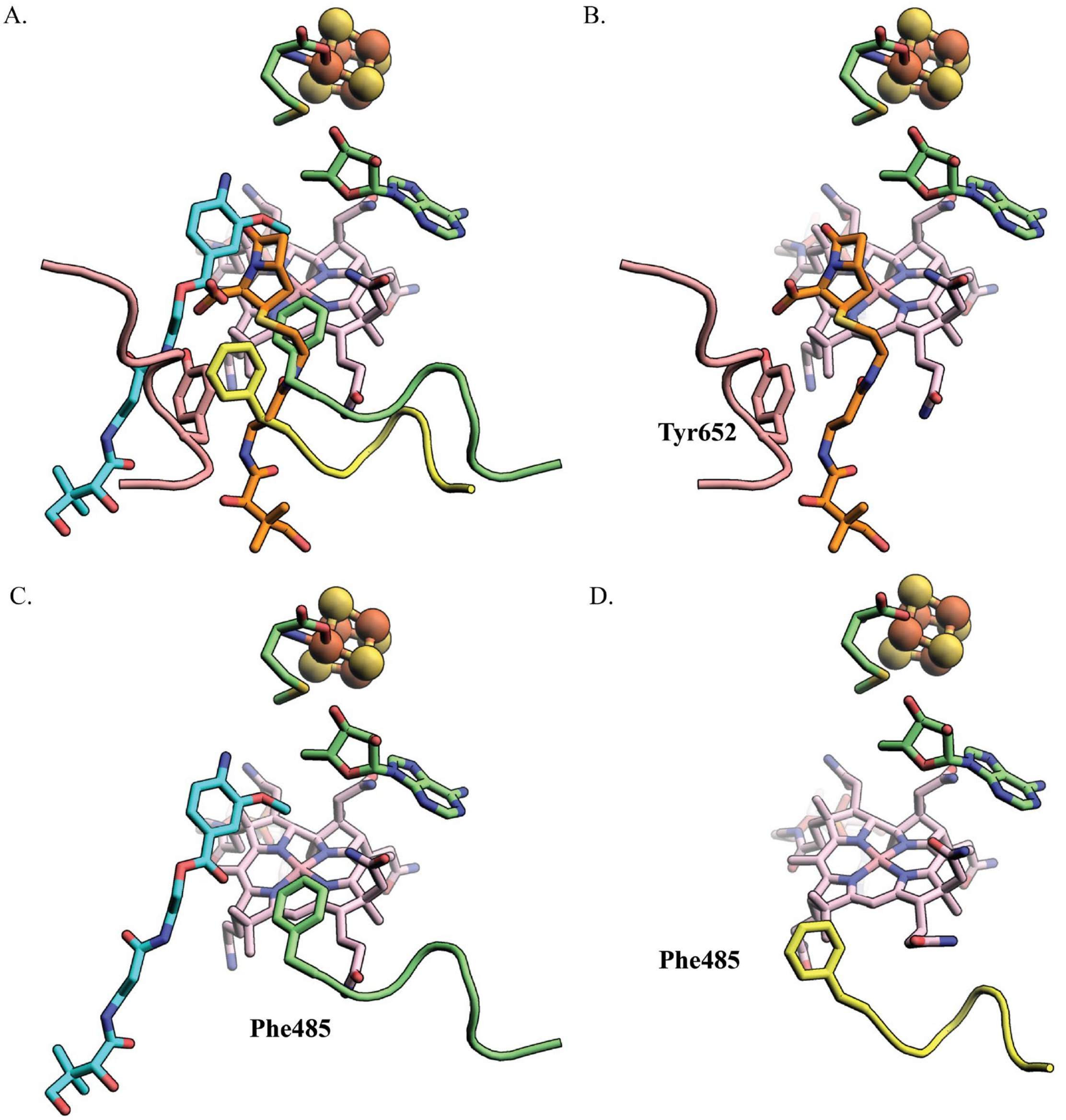


**Figure S12. A.** Overlay of TokK (PDB ID: 7KDY) with *cc*CysS-OMe substrate structure and *cc*CysS-no substrate structure shows Tyr652 of TokK is in a proximal position to Phe485 of *cc*CysS. Tyr652 of TokK and the corresponding loop are shown in pink color. Phe485 of *cc*CysS-OMe substrate structure and the corresponding loop are shown in green color. Phe485 of *cc*CysS-no substrate structure and the corresponding loop are shown in yellow color. The substrate of TokK is shown in orange color, and the substrate of *cc*CysS is shown in cyan color. To make the presentation clear, each overlaid structure is presented individually in Figures S11B (TokK), C (ccCysS-OMe substrate), and D (ccCysS-no substrate), with the same color choices.

A.

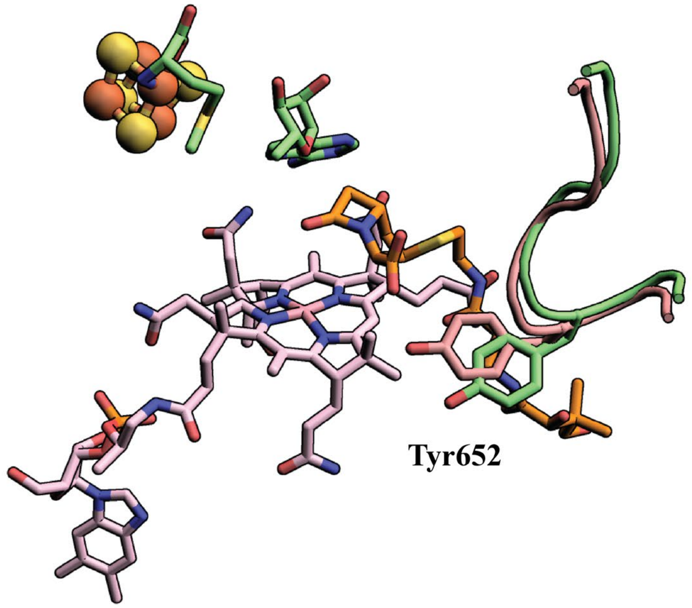


B.

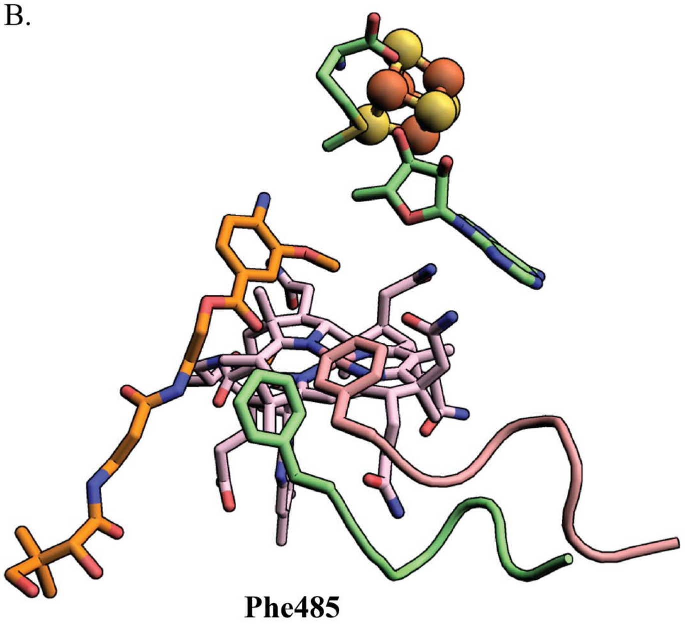


**Figure S13. A.** Overlay of TokK-carbapenem structure (PDB ID: 7KDY) and TokK-no substrate structure (PDB ID: 7KDX). Tyr652 and the corresponding loop in TokK-carbapenem structure (shown in pink) have no significant change compared with those of TokK-no substrate structure (shown in green); **B.** Overlay of *cc*CysS-OMe substrate structure with *cc*CysS-no substrate structure. Phe485 and the corresponding loop in *cc*CysS-OMe substrate structure (shown in pink) has a signification conformational change compared with those of *cc*CysS-no substrate structure (shown in green).

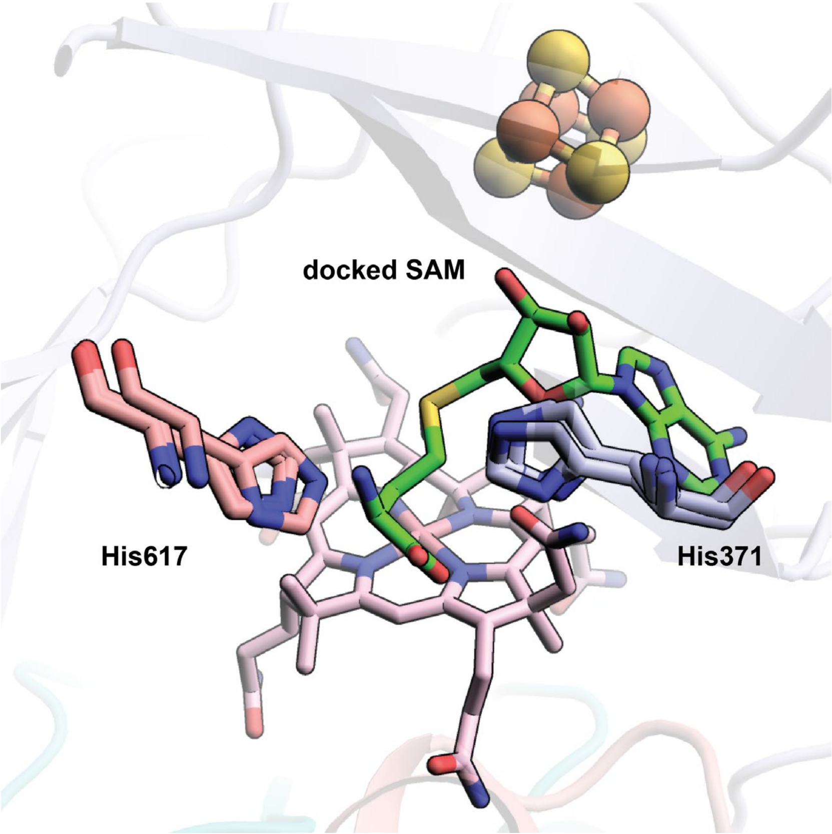


**Figure S14.** Overlaying the SAM-docked *cc*CysS structure with the OMe-bound *cc*CysS structure and the no-substrate-bound *cc*CysS structure revealed that His617 and His371 undergo no major conformational changes to accommodate the interaction with the Met moiety of SAM for SN2 methylation of Cbl.

Sequence of the codon-optimized gene of the *cb*CysS (UniProt ID: A0A0H4NV78) as supplied by Gene Universal:

5′-
CATATGGCAAATCAGCGTGTGGCATTTATTGAACTGACCGTTTTTAGCGGTGT
TTATCCGCTGGCAAGCGGTTATATGCGTGGTGTTGCAGAACAGAATCCGCTG
ATTCGTGAAAGCTGTAGCTTTGAAATTCATAGCATCTGCATTAACGATGATCG
CTTCGAAGATAAGCTGAACAAAATTGATGCAGATGTGTATGCCATTAGCTGC
TATGTTTGGAATATGGGTTTTGTGAAACGTTGGCTGCCGACACTGACCGCACG
TAAACCGAATGCACATATCATTTTAGGTGGTCCGCAGGTTATGAATCATGGTG
CACAGTATCTGGATCCGGGTAATGAACGTGTTGTTCTGTGTAATGGTGAAGG
CGAATATACCTTTGCAAATTATCTGGCAGAACTGTGTAGTCCGCAGCCGGATC
TGGGTAAAGTTAAAGGTCTGAGTTTTATCGCAACGGCGAACTGATTACCAC
CGAACCGCAGGCACGTATTCAGGATCTGAACACCGTTCCGAGTCCGTATCTG
GAAGGTTATTTTGATAGCGAGAAATATGTTTGGGCACCGCTGGAAACCAATC
GTGGTTGTCCGTATCAGTGTACCTATTGTTTTTGGGGTGCAGCAACCAATAGC
CGTGTGTTTAAAAGCGATATGGATCGTGTTAAAGCCGAAATTACCTGGCTGA
GCCAGCATCGTGCCTTTTATATCTTTATTACCGATGCCAATTTTGGCATGCTG
ACCCGTGATATTGAAATTGCACAGCATATTGCCGAATGCAAACGCAAATATG
GCTATCCGCTGACCATTTGGCTGAGTGCAGCCAAAAATAGTCCGGATCGTGT
GACCCAGATTACCCGTATTCTGAGCCAAGAAGGTCTGATTAGCACCCAGCCG
GTTAGCCTGCAGACCATGGATGCAAATACCCTGAAAAGCGTTAAACGCGGTA
ACATTAAAGAAAGCGCATATCTGAGCCTGCAAGAAGAACTGCATCGTAGCAA
ACTGAGCAGCTTTGTTGAAATGATTTGGCCGTTACCGGGTGAAACCCTGGAA
ACCTTTCGTGAAGGTATTGGTAAACTGTGCAGCTATGATGCCGATGCAATTCT
GATTCATCATCTGCTGCTGATTAATAACGTTCCGATGAATAGCCAGCGCGAA
GAATTTAAACTGGAAGTGAGCAATGATGAAGATCCGAATAGCGAAGCACAG
GTTGTTGTTGCAACCAAAGATGTTACCCGTGAAGAATACAAAGAAGGTGTGC
GTTTTGGTTATCATCTGACCAGCCTGTATAGTCTGCGTGCACTGCGTTTTGTTG
GTCGTTATCTGGATAAACAGGGTCGTCTGGCATTTAAAGATCTGATTAGTAGC
TTTAGCGAGTACTGCAAACGTAATCCGGATCATCCGTATACACAGTATATTAC
CAGCGTTATTGATGGCACCAGCCAGAGCAAATTTAGCGCAAATGGTGGTATT
TTTCATGTGACCCTGCATGAATTTCGTCGCGAATTTGATCAGCTGCTGTTTGG
TTTTATTCAGACCCTGGGTATGATGAATGATGAACTGCTGGAATTTCTGTTCG
AAATGGATCTGCTGAATCGTCCGCATGTTTATAGCAATACCCCGATTAACAAT
GGCGAAGGTCTGCTGAAACATGTTACCGTTGTTAGCAAAGAAAAAGATGCCA
TTGTTCTGCGTGTGCCGGAAAAATATGCACAGCTGACCAGCGAACTGTTAGG
TCTGGAAGGCGCACCGAGCACCAGCCTGCGTGTTAAATATCGTGGCACCCAG
ATGCCGTTTATGGCGAATAAACCGTATGAAGATAATCTGAGCTATTGCGAAG
CCAAACTGCATAAAATGGGTAGCATTCTGCCGGTTTGGGAAAGCGCAGTGCC
GAGCCGTACACCGGTTCGTCGTCCTCAGGTTGCAGTTGCAGGTTAACTCGAG-
3′

Sequence of the codon-optimized gene of the *cc*CysS (UniProt ID: A0A3A8HCN5) as supplied by Gene Universal:

5′-
CATATGGGTGCAATGGTTAATCAGCGTGTTGCATTTATTGAACTGACCGTTTT
TGCCGGTGTTTATCCGCTGGCAAGCGGTTATATGCGTGGTGTTGCAGAACAG
AATGCAGCAATTAAAGATGCCTGCAGCTTTGAAATTCATAGCATCTGCATTA
ACGACAACCGTTTTGAAGATCGTCTGAATGCAATTGATGCAGATGTTTATGCC
ATTAGCTGCTATGTTTGGAATATGGGTTTTGTGAAACGTTGGCTGCCGACACT
GACCGCACGTAAACCGCATGCACATGTTATTTTAGGTGGTCCGCAGGTTATG
AATCATGGTGCACGTTATCTGGATCCGGGTAATGAACGTGTTGTTCTGTGTAA
TGGTGAAGGCGAATATACCTTTGCAAATTATCTGGCGGAAATTTGTAGTCCG
GAACCGGATCTGGGTAAAGTTAAAGGCCTGACCTTTTATCGCAATGGTGAAC
TGATTACCAGCGCACCGCAAGAACGTATTCAGGATCTGAATGCCATTCCGAG
TCCGTATCTGGAAGGTTATTTTGATAGCGAGAAATATGTTTGGGCACCGATTG
AAACCAATCGTGGTTGTCCGTATCAGTGTACCTATTGTTTTTGGGGTGCAGCA
ACCAATAGCCGTGTGTTTAAAACCGATATGGATCGTGTTAAAGCCGAAATTA
CCTGGCTGAGCCAGCGTCGTGCCTTTTATATCTTTATTACCGATGCCAATTTTG
GCATGCTGACCCGTGATATTGAAATTGCACAGCATATTGCCGAATGCAAACG
CAAATATGGCTATCCGCTGACCGTGTGGCTGAGTGCAGCCAAAAATAGTCCG
GATCGTGTGACCCAGATTACCCGTATTCTGAGCCAAGAAGGTCTGATTAGCA
CCCAGCCGGTTAGCCTGCAGACCATGGATGCAAATACCCTGAAAAGCGTTAA
ACGCGGTAACATTAAAGAAAGCGCATATCTGAATCTGCAAGAAGAACTGCGT
CGTAGCAAACTGAGCAGCTTTGTTGAAATGATTTGGCCGTTACCGGGTGAAA
CCCTGGAAACCTTTAAAGAGGGTATTGGTAAACTGTGTAGCTATGAAGCAGA
TGCCATTCTGATTCATCATCTGCTGCTGATTAATAACGTTCCGATGAATGCAC
AGCGCGAAGAATTTAATCTGGAAGTGAGCAATGATGAAGATCCGAATAGCG
AAGCACAGGTTGTTGTTGCAACCCGTGATGTTACCCGTGAAGAATACAAAGA
AGGTGTGCGTTTTGGTTATCATCTGACCAGCCTGTATAGTCTGCGTGCACTGC
AGTTTGTTGGTAAATATCTGGATAAACAGGGTCTGCTGGCATTCAAAGATCTG
ATTAGTAGCTTTAGCGATTACTGCAAACGTTTTCCGGATCATCCGTATACACA
GTATATTAGCAGCATTATTGATGGTAGCAGCCAGAGCAAATTTAGCGCAAAT
GGTGGTATTTTTCATGTGACCCTGCATGAATTTCGTCGCGAATTTGATCAGCT
GCTGGCAGGTTTTCTGCAGAGCCTGGGTATGATGCATACCGAACCGCTGGAA
TTTCTGTTTGATCTGGATCTGCTGAATCGTCCGCATGTTTATAGCAATACACC
GGTTACCAATGGTGATGGTCTGCTGAAACATGTTACCGTTGTTGCCAAAGAA
AAAGATGCACTGGTTGTTCATATCCCGGAAAAATATGTTCAGCTGGCATGGG
AAATGCTGCGTCTGGATGGTGCACCGAGCACACGTATGCGTGTTAAATATCG
TGGTGCACAGATGCCGTTTATGGCAAATAAACCGTATGAAGATAACCTGAGC
TATTGCGAAGCAAAACTGCATAAAATGGGTAGCATTCTGCCGGTTTGGGAAC
CTGCAGTTCCGAGCATTGCACCGGTTCGTCGTCCTCAGGTTGCAGTTGCAAGC
TAACTCGAG-3′

Sequence of the codon-optimized gene of the *cc*CysS-W271A as supplied by Gene Universal:

5′-
CATATGGGCGCTATGGTGAATCAGCGCGTGGCATTCATTGAACTGACCGTGTT
TGCCGGTGTTTATCCGCTGGCAAGCGGTTATATGCGTGGTGTTGCAGAACAG
AATGCAGCCATTAAGGATGCCTGCAGCTTTGAAATTCATAGTATTTGTATCAA
CGACAACCGCTTTGAAGATCGTCTGAATGCCATTGATGCAGATGTTTATGCCA
TTAGTTGCTATGTGTGGAATATGGGTTTTGTGAAACGTTGGCTGCCGACCCTG
ACCGCCCGCAAACCGCATGCCCATGTGATTCTGGGTGGCCCGCAGGTTATGA
ATCATGGTGCACGTTATCTGGATCCGGGCAATGAACGTGTTGTTCTGTGTAAT
GGTGAAGGCGAATATACCTTTGCAAATTATCTGGCAGAAATTTGTAGTCCGG
AACCGGATCTGGGTAAAGTTAAAGGTCTGACCTTTTATCGCAATGGCGAACT
GATTACCAGCGCCCCGCAGGAACGTATTCAGGATCTGAATGCCATCCCGAGC
CCGTATCTGGAAGGTTATTTTGATAGTGAAAAGTATGTGTGGGCCCCGATTGA
AACCAATCGCGGCTGTCCGTATCAGTGTACCTATTGCTTTTGGGGCGCCGCCA
CCAATAGCCGTGTGTTTAAAACCGATATGGATCGTGTTAAAGCCGAAATTAC
CTGGCTGAGTCAGCGTCGTGCCTTTTATATTTTTATTACCGATGCAAACTTCG
GTATGCTGACCCGTGATATTGAAATTGCACAGCATATTGCCGAATGTAAACG
TAAATATGGTTATCCGCTGACCGTTGCCCTGAGCGCCGCCAAAAATAGTCCG
GATCGTGTGACCCAGATTACCCGTATTCTGAGCCAGGAAGGTCTGATTAGCA
CCCAGCCGGTTAGTCTGCAGACCATGGATGCAAATACCCTGAAAAGCGTGAA
ACGCGGTAATATTAAGGAAAGTGCCTATCTGAATCTGCAGGAAGAACTGCGC
CGCAGTAAACTGAGTAGCTTTGTTGAAATGATTTGGCCGCTGCCGGGCGAAA
CCCTGGAAACCTTTAAAGAAGGTATTGGCAAACTGTGTAGTTATGAAGCAGA
TGCCATTCTGATTCATCATCTGCTGCTGATTAATAATGTGCCGATGAATGCAC
AGCGTGAAGAATTCAATCTGGAAGTGAGCAATGATGAAGATCCGAATAGCG
AAGCACAGGTTGTTGTTGCAACCCGCGATGTTACCCGTGAAGAATATAAAGA
AGGTGTTCGTTTTGGTTATCATCTGACCAGTCTGTATAGCCTGCGTGCCCTGC
AGTTTGTGGGTAAATATCTGGATAAACAGGGTCTGCTGGCCTTTAAAGATCTG
ATTAGCAGTTTTAGCGATTATTGTAAACGTTTTCCGGATCATCCGTATACCCA
GTATATTAGCAGCATTATTGATGGTAGTAGCCAGAGTAAATTTTCAGCCAATG
GCGGCATTTTTCATGTGACCCTGCATGAATTTCGTCGCGAATTTGATCAGCTG
CTGGCAGGTTTTCTGCAGAGCCTGGGCATGATGCATACCGAACCGCTGGAAT
TTCTGTTTGATCTGGATCTGCTGAATCGTCCGCATGTGTATAGTAATACCCCG
GTTACCAATGGCGATGGTCTGCTGAAACATGTTACCGTTGTTGCCAAAGAAA
AAGATGCCCTGGTGGTGCATATTCCGGAAAAATATGTGCAGCTGGCCTGGGA
AATGCTGCGTCTGGATGGCGCACCGAGCACCCGTATGCGTGTTAAATATCGT
GGTGCCCAGATGCCGTTTATGGCAAATAAGCCGTATGAAGATAATCTGAGCT
ATTGTGAAGCCAAACTGCATAAAATGGGTAGCATTCTGCCGGTTTGGGAACC
GGCCGTTCCGAGCATTGCACCGGTTCGTCGTCCGCAGGTTGCCGTGGCCAGTT
AACTCGAG -3′

Sequence of the codon-optimized gene of the *cc*CysS-W271H as supplied by Gene Universal:

5′-
CATATGGGTGCAATGGTGAATCAGCGTGTGGCCTTTATTGAACTGACCGTTTT
TGCCGGCGTTTATCCGCTGGCCAGTGGCTATATGCGTGGTGTGGCCGAACAG
AATGCAGCAATTAAGGATGCATGCAGTTTTGAAATTCATAGTATTTGTATCAA
CGACAACCGTTTTGAAGATCGTCTGAATGCCATTGATGCCGATGTGTATGCCA
TTAGCTGTTATGTTTGGAATATGGGTTTTGTTAAGCGTTGGCTGCCGACCCTG
ACCGCCCGCAAACCGCATGCCCATGTTATTCTGGGCGGCCCGCAGGTTATGA
ATCATGGTGCACGCTATCTGGATCCGGGTAATGAACGCGTTGTGCTGTGTAAT
GGTGAAGGCGAATATACCTTTGCCAATTATCTGGCCGAAATTTGCAGTCCGG
AACCGGATCTGGGTAAAGTGAAAGGCCTGACCTTTTATCGTAATGGCGAACT
GATTACCAGCGCACCGCAGGAACGCATTCAGGATCTGAATGCAATTCCGAGC
CCGTATCTGGAAGGCTATTTTGATAGCGAAAAATATGTTTGGGCACCGATTG
AAACCAATCGCGGTTGCCCGTATCAGTGCACCTATTGTTTTTGGGGTGCCGCA
ACCAATAGTCGCGTTTTTAAAACCGATATGGATCGTGTGAAAGCAGAAATTA
CCTGGCTGAGTCAGCGTCGCGCATTTTATATTTTTATTACCGATGCAAACTTC
GGCATGCTGACCCGCGATATTGAAATTGCCCAGCATATTGCAGAATGCAAAC
GCAAATATGGTTATCCGCTGACCGTGCATCTGAGCGCCGCAAAAAATAGCCC
GGATCGCGTGACCCAGATTACCCGCATTCTGAGTCAGGAAGGTCTGATTAGC
ACCCAGCCGGTTAGTCTGCAGACCATGGATGCAAATACCCTGAAAAGCGTGA
AACGTGGTAATATTAAGGAAAGCGCCTATCTGAATCTGCAGGAAGAACTGCG
CCGTAGTAAACTGAGCAGTTTTGTTGAAATGATTTGGCCGCTGCCGGGTGAA
ACCCTGGAAACCTTTAAAGAAGGCATTGGTAAACTGTGCAGCTATGAAGCAG
ATGCCATTCTGATTCATCATCTGCTGCTGATTAATAATGTGCCGATGAATGCC
CAGCGCGAAGAATTCAATCTGGAAGTTAGTAATGATGAAGATCCGAATAGTG
AAGCACAGGTTGTGGTGGCAACCCGTGATGTTACCCGCGAAGAATATAAAGA
AGGCGTTCGTTTTGGTTATCATCTGACCAGTCTGTATAGCCTGCGTGCACTGC
AGTTTGTTGGTAAATATCTGGATAAACAGGGCCTGCTGGCATTCAAAGATCT
GATTAGTAGCTTTAGCGATTATTGCAAACGCTTTCCGGATCATCCGTATACCC
AGTATATTAGTAGTATTATTGACGGTAGCAGCCAGAGCAAATTTTCTGCAAAT
GGTGGTATTTTTCACGTTACCCTGCATGAATTTCGCCGCGAATTTGATCAGCT
GCTGGCCGGTTTTCTGCAGAGTCTGGGTATGATGCATACCGAACCGCTGGAA
TTTCTGTTTGATCTGGATCTGCTGAATCGTCCGCATGTGTATAGCAATACCCC
GGTGACCAATGGTGACGGTCTGCTGAAACATGTGACCGTGGTTGCAAAAGAA
AAAGATGCCCTGGTTGTGCATATTCCGGAAAAATATGTGCAGCTGGCATGGG
AAATGCTGCGTCTGGATGGTGCCCCGAGCACCCGTATGCGCGTTAAATATCG
TGGTGCCCAGATGCCGTTTATGGCAAATAAGCCGTATGAAGATAATCTGAGT
TATTGCGAAGCAAAACTGCATAAAATGGGCAGCATTCTGCCGGTGTGGGAAC
CGGCCGTTCCGAGCATTGCCCCGGTTCGTCGTCCGCAGGTTGCAGTGGCCAG
CTAACTCGAG -3′

Sequence of the codon-optimized gene of the *cc*CysS-W271Q as supplied by Gene Universal:

5′-
CATATGGGCGCTATGGTGAATCAGCGTGTTGCATTCATTGAACTGACCGTGTT
TGCCGGTGTTTATCCGCTGGCCAGTGGTTATATGCGTGGCGTTGCCGAACAGA
ATGCAGCCATTAAGGATGCATGCAGTTTTGAAATTCATAGCATTTGCATTAAC
GACAATCGCTTTGAAGATCGCCTGAATGCCATTGATGCCGATGTTTATGCAAT
TAGTTGCTATGTTTGGAATATGGGCTTTGTTAAACGTTGGCTGCCGACCCTGA
CCGCCCGTAAACCGCATGCCCATGTTATTCTGGGTGGTCCGCAGGTTATGAAT
CATGGTGCCCGCTATCTGGATCCGGGCAATGAACGTGTTGTTCTGTGTAATGG
TGAAGGCGAATATACCTTTGCAAATTATCTGGCCGAAATTTGTAGCCCGGAA
CCGGATCTGGGTAAAGTTAAAGGTCTGACCTTTTATCGCAATGGCGAACTGA
TTACCAGCGCACCGCAGGAACGCATTCAGGATCTGAATGCAATTCCGAGTCC
GTATCTGGAAGGCTATTTTGATAGCGAAAAATATGTTTGGGCCCCGATTGAA
ACCAATCGTGGCTGTCCGTATCAGTGTACCTATTGCTTTTGGGGTGCAGCAAC
CAATAGCCGCGTTTTTAAAACCGATATGGATCGTGTTAAAGCAGAAATTACC
TGGCTGAGCCAGCGCCGCGCCTTTTATATTTTTATTACCGATGCCAATTTCGG
TATGCTGACCCGTGATATTGAAATTGCCCAGCATATTGCCGAATGTAAACGTA
AATATGGTTATCCGCTGACCGTGCAGCTGAGTGCCGCCAAAAATAGTCCGGA
TCGTGTTACCCAGATTACCCGTATTCTGAGTCAGGAAGGCCTGATTAGTACCC
AGCCGGTTAGTCTGCAGACCATGGATGCAAATACCCTGAAAAGCGTTAAACG
TGGCAATATTAAGGAAAGTGCCTATCTGAATCTGCAGGAAGAACTGCGTCGT
AGTAAACTGAGTAGCTTTGTTGAAATGATTTGGCCGCTGCCGGGCGAAACCC
TGGAAACCTTTAAAGAAGGTATTGGTAAACTGTGTAGCTATGAAGCAGATGC
AATTCTGATTCATCATCTGCTGCTGATTAATAATGTTCCGATGAATGCACAGC
GTGAAGAATTCAATCTGGAAGTTAGCAATGATGAAGATCCGAATAGTGAAGC
ACAGGTTGTTGTTGCCACCCGTGATGTTACCCGTGAAGAATATAAAGAAGGT
GTTCGTTTTGGTTATCATCTGACCAGTCTGTATAGTCTGCGCGCACTGCAGTTT
GTTGGTAAATATCTGGATAAACAGGGTCTGCTGGCATTCAAAGATCTGATTA
GTAGTTTTAGCGATTACTGTAAACGCTTTCCGGATCATCCGTATACCCAGTAT
ATTAGCAGTATTATTGACGGTAGTAGCCAGAGTAAATTTTCTGCAAATGGCG
GTATTTTTCATGTTACCCTGCATGAATTTCGTCGCGAATTTGATCAGCTGCTG
GCAGGCTTTCTGCAGAGTCTGGGTATGATGCATACCGAACCGCTGGAATTTCT
GTTTGATCTGGATCTGCTGAATCGTCCGCATGTTTATAGCAATACCCCGGTTA
CCAATGGTGACGGCCTGCTGAAACATGTTACCGTTGTTGCCAAAGAAAAAGA
TGCACTGGTTGTGCATATTCCGGAAAAATATGTGCAGCTGGCCTGGGAAATG
CTGCGCCTGGATGGCGCCCCGAGCACCAGAATGCGTGTTAAATATCGCGGTG
CACAGATGCCGTTTATGGCAAATAAGCCGTATGAAGATAATCTGAGTTATTG
CGAAGCCAAACTGCATAAAATGGGTAGTATTCTGCCGGTTTGGGAACCGGCC
GTGCCGAGTATTGCCCCGGTGCGTCGCCCGCAGGTTGCCGTGGCAAGCTAAC
TCGAG -3′

Sequence of the codon-optimized gene of the *cc*CysS-F485W as supplied by Gene Universal:

5′-
CATATGGGTGCAATGGTGAATCAGCGTGTTGCCTTTATTGAACTGACCGTGTT
TGCCGGCGTGTATCCGCTGGCCAGTGGCTATATGCGCGGCGTGGCCGAACAG
AATGCCGCCATTAAGGATGCCTGCAGTTTTGAAATTCATAGCATTTGCATTAA
CGACAATCGTTTTGAAGATCGTCTGAATGCCATTGATGCCGATGTGTATGCCA
TTAGTTGCTATGTTTGGAATATGGGTTTTGTTAAACGCTGGCTGCCGACCCTG
ACCGCACGCAAACCGCATGCCCATGTGATTCTGGGTGGCCCGCAGGTTATGA
ATCATGGTGCCCGCTATCTGGATCCGGGCAATGAACGCGTTGTGCTGTGTAAT
GGTGAAGGCGAATATACCTTTGCCAATTATCTGGCAGAAATTTGTAGCCCGG
AACCGGATCTGGGTAAAGTTAAAGGCCTGACCTTTTATCGCAATGGTGAACT
GATTACCAGCGCCCCGCAGGAACGTATTCAGGATCTGAATGCCATCCCGAGT
CCGTATCTGGAAGGCTATTTTGATAGCGAAAAATATGTGTGGGCACCGATTG
AAACCAATCGTGGCTGTCCGTATCAGTGCACCTATTGTTTTTGGGGCGCCGCA
ACCAATAGTCGTGTTTTTAAAACCGATATGGATCGCGTGAAAGCAGAAATTA
CCTGGCTGAGCCAGCGTCGTGCCTTTTATATTTTTATTACCGATGCAAACTTC
GGTATGCTGACCCGTGATATTGAAATTGCACAGCATATTGCAGAATGTAAAC
GTAAATATGGTTACCCGCTGACCGTTTGGCTGAGCGCCGCCAAAAATAGCCC
GGATCGTGTGACCCAGATTACCCGTATTCTGAGCCAGGAAGGTCTGATTAGT
ACCCAGCCGGTTAGTCTGCAGACCATGGATGCAAATACCCTGAAAAGCGTTA
AACGCGGTAATATTAAGGAAAGCGCATATCTGAATCTGCAGGAAGAACTGCG
TCGCAGTAAACTGAGCAGCTTTGTGGAAATGATTTGGCCGCTGCCGGGCGAA
ACCCTGGAAACCTTTAAAGAAGGCATTGGTAAACTGTGCAGTTATGAAGCCG
ATGCCATTCTGATTCATCATCTGCTGCTGATTAATAATGTGCCGATGAATGCC
CAGCGTGAAGAATTCAATCTGGAAGTTAGCAATGATGAAGATCCGAATAGCG
AAGCACAGGTGGTTGTTGCAACCCGCGATGTTACCCGCGAAGAATATAAAGA
AGGTGTGCGCTTTGGTTATCATCTGACCAGCCTGTATAGCCTGCGTGCCCTGC
AGTTTGTGGGCAAATATCTGGATAAACAGGGCCTGCTGGCCTTTAAAGATCT
GATTAGCAGCTTTAGCGATTATTGCAAACGTTTTCCGGATCATCCGTATACCC
AGTATATTAGTAGCATTATTGACGGCAGTAGTCAGAGCAAATGGAGTGCCAA
TGGTGGCATTTTTCATGTTACCCTGCATGAATTTCGCCGTGAATTTGATCAGC
TGCTGGCAGGTTTTCTGCAGAGTCTGGGTATGATGCATACCGAACCGCTGGA
ATTTCTGTTTGATCTGGATCTGCTGAATCGCCCGCATGTGTATAGCAATACCC
CGGTGACCAATGGCGATGGCCTGCTGAAACATGTGACCGTGGTTGCAAAAGA
AAAAGATGCACTGGTGGTGCATATTCCGGAAAAATATGTTCAGCTGGCCTGG
GAAATGCTGCGCCTGGATGGTGCACCGAGTACCCGCATGCGTGTGAAATATC
GTGGTGCACAGATGCCGTTTATGGCCAATAAGCCGTATGAAGATAATCTGAG
TTATTGTGAAGCCAAACTGCATAAAATGGGTAGCATTCTGCCGGTTTGGGAA
CCGGCAGTTCCGAGTATTGCCCCGGTTCGCCGCCCGCAGGTGGCAGTGGCAA
GCTAACTCGAG -3′

Sequence of the codon-optimized gene of *cc*CysS-F485Y as supplied by Gene Universal:

5′-
CATATGGGTGCAATGGTGAATCAGCGCGTGGCCTTTATTGAACTGACCGTGTT
TGCAGGCGTTTATCCGCTGGCCAGCGGCTATATGCGTGGTGTGGCAGAACAG
AATGCCGCCATTAAGGATGCCTGTAGCTTTGAAATTCATAGCATTTGTATCAA
CGACAATCGTTTTGAAGATCGCCTGAATGCCATTGATGCAGATGTGTATGCA
ATTAGTTGCTATGTTTGGAATATGGGTTTTGTGAAACGTT
GGCTGCCGACCCTGACCGCCCGCAAACCGCATGCACATGTGATTCTGGGCGG
TCCGCAGGTTATGAATCATGGCGCCCGTTATCTGGATCCGGGCAATGAACGC
GTTGTTCTGTGTAATGGCGAAGGTGAATATACCTTTGCCAATTATCTGGCCGA
AATTTGCAGTCCGGAACCGGATCTGGGCAAAGTGAAAGGTCTGACCTTTTAT
CGCAATGGTGAACTGATTACCAGTGCACCGCAGGAACGTATTCAGGATCTGA
ATGCCATCCCGAGTCCGTATCTGGAAGGTTATTTTGATAGCGAAAAATATGTG
TGGGCACCGATTGAAACCAATCGCGGTTGCCCGTATCAGTGTACCTATTGCTT
TTGGGGTGCAGCCACCAATAGCCGTGTGTTTAAAACCGATATGGATCGCGTG
AAAGCCGAAATTACCTGGCTGAGTCAGCGTCGTGCCTTTTATATTTTTATTAC
CGATGCAAACTTCGGCATGCTGACCCGTGATATTGAAATTGCACAGCATATT
GCAGAATGCAAACGCAAATATGGTTATCCGCTGACCGTTTGGCTGAGCGCAG
CAAAAAATAGTCCGGATCGTGTGACCCAGATTACCCGCATTCTGAGTCAGGA
AGGCCTGATTAGCACCCAGCCGGTTAGCCTGCAGACCATGGATGCAAATACC
CTGAAAAGCGTGAAACGCGGCAATATTAAGGAAAGTGCCTATCTGAATCTGC
AGGAAGAACTGCGCCGTAGCAAACTGAGTAGCTTTGTTGAAATGATTTGGCC
GCTGCCGGGCGAAACCCTGGAAACCTTTAAAGAAGGTATTGGTAAACTGTGC
AGCTATGAAGCAGATGCAATTCTGATTCATCATCTGCTGCTGATTAATAATGT
TCCGATGAATGCACAGCGTGAAGAATTCAATCTGGAAGTTAGCAATGATGAA
GATCCGAATAGCGAAGCCCAGGTTGTTGTTGCAACCCGCGATGTTACCCGCG
AAGAATATAAAGAAGGTGTGCGTTTTGGTTATCATCTGACCAGTCTGTATAGT
CTGCGCGCCCTGCAGTTTGTGGGCAAATATCTGGATAAACAGGGTCTGCTGG
CCTTTAAAGATCTGATTAGTAGTTTTAGTGACTACTGTAAACGCTTTCCGGAT
CATCCGTATACCCAGTATATTAGCAGCATTATTGATGGCAGCAGCCAGAGCA
AATATAGTGCAAATGGCGGTATTTTTCATGTGACCCTGCATGAATTTCGCCGC
GAATTTGATCAGCTGCTGGCCGGTTTTCTGCAGAGCCTGGGCATGATGCATAC
CGAACCGCTGGAATTTCTGTTTGATCTGGATCTGCTGAATCGCCCGCATGTGT
ATAGTAATACCCCGGTTACCAATGGTGACGGTCTGCTGAAACATGTGACCGT
TGTTGCCAAAGAAAAAGATGCACTGGTGGTTCATATTCCGGAAAAATATGTT
CAGCTGGCCTGGGAAATGCTGCGCCTGGATGGTGCCCCGAGTACCCGTATGC
GCGTGAAATATCGCGGTGCACAGATGCCGTTTATGGCAAATAAGCCGTATGA
AGATAATCTGAGTTATTGCGAAGCCAAACTGCATAAAATGGGCAGCATTCTG
CCGGTTTGGGAACCGGCCGTTCCGAGTATTGCCCCGGTGCGTCGCCCGCAGG
TTGCAGTTGCCAGTTAACTCGAG-3′

Sequence of the codon-optimized gene of *cc*CysS-F485L as supplied by Gene Universal:

5′-
CATATGGGTGCTATGGTTAATCAGCGCGTTGCCTTTATTGAACTGACCGTTTT
TGCAGGTGTGTATCCGCTGGCAAGTGGCTATATGCGTGGTGTTGCCGAACAG
AATGCCGCAATTAAGGATGCCTGTAGTTTTGAAATTCATAGCATTTGCATCAA
CGATAATCGTTTTGAAGATCGTCTGAATGCCATTGATGCAGATGTTTATGCCA
TTAGCTGTTATGTTTGGAATATGGGCTTTGTTAAACGTTGGCTGCCGACCCTG
ACCGCCCGCAAACCGCATGCCCATGTGATTCTGGGTGGCCCGCAGGTTATGA
ATCATGGTGCCCGCTATCTGGATCCGGGTAATGAACGCGTTGTTCTGTGCAAT
GGCGAAGGTGAATATACCTTTGCAAATTATCTGGCAGAAATTTGTAGCCCGG
AACCGGATCTGGGTAAAGTGAAAGGTCTGACCTTTTATCGTAATGGCGAACT
GATTACCAGCGCCCCGCAGGAACGCATTCAGGATCTGAATGCAATTCCGAGT
CCGTATCTGGAAGGTTATTTTGATAGTGAAAAGTATGTGTGGGCCCCGATTGA
AACCAATCGTGGTTGTCCGTATCAGTGTACCTATTGCTTTTGGGGCGCCGCAA
CCAATAGTCGCGTTTTTAAAACCGATATGGATCGTGTGAAAGCAGAAATTAC
CTGGCTGAGCCAGCGTCGTGCATTTTATATTTTTATTACCGATGCAAACTTCG
GTATGCTGACCCGTGATATTGAAATTGCCCAGCATATTGCCGAATGCAAACG
CAAATATGGTTATCCGCTGACCGTTTGGCTGAGTGCCGCAAAAAATAGTCCG
GATCGCGTTACCCAGATTACCCGTATTCTGAGTCAGGAAGGTCTGATTAGTAC
CCAGCCGGTTAGTCTGCAGACCATGGATGCAAATACCCTGAAAAGTGTGAAA
CGCGGTAATATTAAGGAAAGTGCCTATCTGAATCTGCAGGAAGAACTGCGTC
GCAGTAAACTGAGTAGCTTTGTTGAAATGATTTGGCCGCTGCCGGGTGAAAC
CCTGGAAACCTTTAAAGAAGGTATTGGCAAACTGTGCAGTTATGAAGCAGAT
GCCATTCTGATTCATCATCTGCTGCTGATTAATAATGTTCCGATGAATGCCCA
GCGTGAAGAGTTTAATCTGGAAGTGAGTAATGATGAAGATCCGAATAGTGAA
GCCCAGGTGGTTGTTGCCACCCGCGATGTGACCCGTGAAGAATATAAAGAAG
GTGTGCGTTTTGGTTATCATCTGACCAGCCTGTATAGCCTGCGTGCCCTGCAG
TTTGTTGGCAAATATCTGGATAAACAGGGTCTGCTGGCATTCAAAGATCTGAT
TAGTAGCTTTAGTGATTACTGCAAACGTTTTCCGGATCATCCGTATACCCAGT
ATATTAGCAGTAT
TATTGACGGTAGTAGCCAGAGTAAACTGAGCGCAAATGGCGGCATTTTTCAT
GTTACCCTGCATGAATTTCGTCGTGAATTTGATCAGCTGCTGGCAGGTTTTCT
GCAGAGCCTGGGCATGATGCATACCGAACCGCTGGAATTTCTGTTTGATCTG
GATCTGCTGAATCGCCCGCATGTGTATAGCAATACCCCGGTTACCAATGGTG
ACGGTCTGCTGAAACATGTGACCGTTGTGGCCAAAGAAAAAGATGCCCTGGT
TGTTCATATTCCGGAAAAATATGTGCAGCTGGCCTGGGAAATGCTGCGTCTG
GATGGTGCCCCGAGCACCCGTATGCGCGTTAAATATCGTGGTGCACAGATGC
CGTTTATGGCAAATAAGCCGTATGAAGATAATCTGAGTTATTGTGAAGCCAA
ACTGCATAAAATGGGTAGTATTCTGCCGGTGTGGGAACCGGCAGTTCCGAGC
ATTGCACCGGTGCGTCGTCCGCAGGTTGCAGTTGCCAGTTAACTCGAG-3′

**Table S1**. Crystallographic data table for *cc*CysS structures obtained.

| | *cc*CysS + OMe | *cc*CysS + OEt | *cc*CysS no substrate |
|---|---|---|---|
| **Data collection** | | | |
| Wavelength | 0.97872 | 1.03317 | 1.03580 |
| Resolution range* | 50 – 1.75 (1.78 - 1.75) | 50 – 2.00 (2.03 – 2.00) | 50 – 1.95 (1.98 – 1.95) |
| Space group | $P\ 1\ 2_1\ 1$ | $P\ 1\ 2_1\ 1$ | $P\ 1\ 2_1\ 1$ |
| Cell dimentions | | | |
| *a, b, c* (Å) | 56.04, 75.29, 77.02 | 56.45, 75.99, 77.05 | 59.28, 77.82, 77.04 |
| α, β, γ (°) | 90, 100.8, 90 | 90, 101.3, 90 | 90, 110.9, 90 |
| $R_{sym}$ or $R_{merge}$ * | 0.070 (0.513) | 0.050 (0.553) | 0.071 (0.522) |
| $R_{pim}$* | 0.027 (0.197) | 0.024 (0.257) | 0.045 (0.360) |
| I / σ * | 27.1 (3.6) | 29.8 (2.9) | 17.4 (3.6) |
| Wilson B-factor | 18.2 | 24.3 | 17.2 |
| $CC_{1/2}$ * | 0.997 (0.890) | 0.998 (0.829) | 0.978 (0.753) |
| Completeness (%) * | 99.1 (98.3) | 98.9 (99.2) | 99.9 (99.9) |
| Redundancy* | 7.7 (7.7) | 5.2 (5.3) | 3.4 (3.0) |
| **Refinement** | | | |
| Resolution (Å) | 1.75 | 2.00 | 1.95 |
| Reflections used in refinement | 62662 | 42349 | 47775 |
| Reflections in $R_{free}$ set (%) | 60303 (5%) | 41182 (4.72%) | 46492 (4.23%) |
| $R_{work}$ / $R_{free}$ | 0.15 / 0.18 | 0.17 / 0.20 | 0.16 / 0.20 |
| Number of non-hydrogen atoms | 5857 | 5622 | 5783 |
| Macromolecules | 5081 | 5063 | 5117 |
| OH-cobalamin | 91 | 91 | 91 |
| Fe/S cluster | 8 | 8 | 8 |
| 5′-dA | 18 | 18 | 18 |
| Met | 9 | 9 | 9 |
| Magnesium-1 | 1 | N/A | N/A |
| Magnesium-2 | 1 | N/A | N/A |
| OMe substrate | 58 | N/A | N/A |
| sodium | N/A | 1 | 1 |
| OEt substrate | N/A | 30 | N/A |
| water | 599 | 411 | 548 |
| *B*-factors (Å$^2$) | | | |
| Macromolecules | 25.6 | 38.43 | 23.75 |
| OH-cobalamin | 14.3 | 25.1 | 14.3 |
| Fe/S cluster | 10.5 | 18.5 | 9.6 |
| 5′-dA | 12.9 | 17.8 | 11.7 |
| Met | 11.5 | 18.2 | 9.6 |
| Magnesium-1 | 18.4 | N/A | N/A |
| Magnesium-2 | 18.5 | N/A | N/A |
| OMe substrate | 25.4 | N/A | N/A |
| sodium | N/A | 22.7 | 16.6 |
| OEt substrate | N/A | 42.6 | N/A |
| water | 32.4 | 35.2 | 29.5 |
| RMS deviations | | | |
| Bond lengths (Å) | 0.009 | 0.007 | 0.008 |
| Bond angles (°) | 1.28 | 1.14 | 1.19 |
| Clashs score | 1.73 | 2.04 | 2.59 |
| Rotamer outliers (%) | 0.18 | 0.18 | 0.18 |
| Ramachandran | | | |
| Most favored (%) | 97.6 | 97.3 | 97.4 |
| Allowed (%) | 2.2 | 2.5 | 2.6 |
| Outliers (%) | 0.2 | 0.2 | 0 |
| Number of TLS groups | 5 | 5 | 5 |
| PDB accession code | 9N1B | 9N1C | 9N1D |

All datasets result from a single protein crystal.
*Values in parentheses are for the highest resolution shell.
